# Single-cell bisulfite-free 5mC and 5hmC sequencing with high sensitivity and scalability

**DOI:** 10.1101/2023.10.04.560836

**Authors:** Yunlong Cao, Yali Bai, Tianjiao Yuan, Liyang Song, Yu Fan, Liuhao Ren, Weiliang Song, Jiahui Peng, Ran An, Qingqing Gu, Yinghui Zheng, Xiaoliang Sunney Xie

## Abstract

Existing single cell bisulfite-based DNA methylation analysis is limited by low DNA recovery, and the measurement of 5hmC at single-base resolution remains challenging. Here we present a bisulfite-free single-cell whole-genome 5mC and 5hmC profiling technique, named Cabernet, which can characterize 5mC and 5hmC at single-base resolution with high genomic coverage. Cabernet utilizes Tn5 transposome for DNA fragmentation, which enables the discrimination between different alleles for measuring hemi-methylation status. Using Cabernet, we revealed the 5mC, hemi-5mC and 5hmC dynamics during early mouse embryo development, uncovering genomic regions exclusively governed by active or passive demethylation. We show that hemi-methylation status can be used to distinguish between pre– and post-replication cells, enabling more efficient cell grouping when integrated with 5mC profile. The property of Tn5 naturally enables Cabernet to achieve high-throughput single-cell methylome profiling, where we probed mouse cortical neurons and embryonic day 7.5 (E7.5) embryos, and constructed the library for thousands of single cells at high efficiency, demonstrating its potential for analyzing complex tissues at substantially low cost. Together, we present a new way of high-throughput methylome and hydroxymethylome detection at single-cell resolution, enabling efficient analysis of epigenetic status of biological systems with complicated nature such as neurons and cancer cells.

**Significance Statement:** Most of current methylation profiling techniques rely on bisulfite treatment, which suffers low DNA recovery. The technique proposed in this study, named Cabernet, can be used to measure 5mC and 5hmC at single-base resolution with high genomic coverage. By using Tn5 transposome, hemi-methylation status can be measured and high-throughput methylome profiling can be achieved. Together, it provides an efficient way to analyze the epigenetic landscape of complicated biological systems.

## Introduction

In mammals, DNA methylation mostly occurs at CpG dinucleotides in the form of 5–methylcytosine (5mC), which plays a vital role in gene regulation, cellular development and disease formation (1, 2). Single-cell DNA methylation sequencing is required to reveal cellular heterogeneity and to study rare samples such as early embryonic development (3–7). However, bisulfite sequencing (BS-seq), the gold standard for mapping mammalian DNA methylation (8, 9), involves harsh chemical reaction that degrades most of the double-stranded DNA (dsDNA), resulting in considerable loss of information (10). Despite huge DNA loss, bisulfite-dependent single-cell methylation sequencing has already been deployed in biological research. For example, a single-cell methylome sequencing method based on reduced representation bisulfite sequencing (RRBS) was developed and applied to probe the methylation in early mouse embryos (11). Yet, this method preferably covers regions with high CpG density and yields a genomic coverage of less 5%. To achieve whole-genome methylation sequencing, other methods such as scBS-seq and scWGBS have been developed based on the post-bisulfite adaptor tagging (PBAT) strategy through random-primed DNA synthesis (12–15). However, multiple rounds of random priming, which are intended to achieve maximum dsDNA recovery, could introduce primer dimers and concatemers which lead to low mapping efficiency. The later reported snmC-seq and snmC-seq2 methods improved read mapping by combining random priming with adaptase reaction (16, 17). Nonetheless, these methods still all involve harsh bisulfite treatment which degrades most of the DNA and limits the overall genomic coverage. An efficient whole-genome methylation detection method at single-cell and single-base resolution is highly preferred. However, to develop such a method, several problems need to be solved.

First, DNA damage caused by bisulfite treatment should be avoided, to which end several bisulfite-free methylome sequencing techniques have been proposed. TET-assisted pyridine borane sequencing (TAPS) uses TET oxidation coupled with pyridine borane reduction to convert modified cytosines to dihydrouracil (DHU) (18). Enzymatic methyl-seq (EM-seq) combines TET2 and BGT protection of 5mC and 5-hydroxymethylcytosine (5hmC), with APOBEC deamination of C to U (19). While these methods allow for better DNA preservation by avoiding bisulfite treatment, the multiple purification steps involved in these methods lead to substantial loss of DNA during the process, limiting their application to single-cell scenarios. Similarly, techniques developed for 5hmC profiling such as 5hmC–DIP and hMe–Seal (20, 21) which are based on affinity enrichment, cannot achieve single-base resolution. The later reported TAB–seq, oxBS-seq and ACE-seq methods, all require high DNA input due to complicated workflow and multiple purification steps, limiting their application to single cells (22–25). To reveal 5mC and 5hmC heterogeneity in cell populations across the entire genome, a bisulfite-free DNA modification profiling approach with minimal DNA loss at single-cell level and single-base resolution is preferred.

Second, characterizing cell-type-specific methylation status in complex tissues requires a scalable single cell-based sequencing technique. Currently established single-cell sequencing methods (12–17), where each single cell is processed in an individual preparation, are highly labor-intensive. Although the application of brute-force strategy can help scale up to a certain extent, as in snmC-seq (16), it’s still a laborious method since every cell produced requires an individual reaction. On the other hand, sci-MET solves this problem by using multiple stages of indexing to label thousands of single cells for pooled library preparation, enabling high-throughput 5mC profiling (26). The employment of barcoded Tn5 transposome during combinatorial indexing is a critical step for the high-throughput of this method; however, the use of bisulfite treatment limits the role of Tn5, necessitating subsequent random-priming. Yet, despite high scalability, this method yields substantially low genomic coverage due to bisulfite conversion. Therefore, a high-throughput single-cell methylation sequencing method with improved DNA coverage is in need.

To address these challenges, we developed a method named Cabernet (Carrier-Assisted Base-conversion by Enzymatic ReactioN with End-Tagging), based on EM-seq. Instead of bisulfite treatment, Cabernet relies on enzymatic conversion where 5mC and 5hmC are converted to 5gmC through TET oxidation and BGT-mediated glycosylation while C is converted to uracil through APOBEC-mediated deamination, following EM-seq protocol (19). Importantly, Cabernet utilizes carrier DNA to minimize DNA loss during purification and achieves scalability through the employment of Tn5 transposome. Together, this technique enables high-throughput detection of whole methylome and hydroxymethylome at single-cell and single-base resolution with high genomic coverage.

## Results

### Validation of Cabernet

The process of Cabernet includes the whole methylation conversion protocol of EM-seq (19), which combines TET-mediated oxidation, BGT-mediated glycosylation and APOBEC-mediated deamination reactions. Given that the original protocol is not applicable to single-cell settings due to huge DNA loss, the following adjustments are made. First, Tn5 insertion, instead of ligation, is used during fragmentation to minimize DNA loss from ultrasonic shearing. Second, carrier dsDNA is introduced to prevent the loss of dsDNA from irretrievable adhesion during purification. Third, library amplification is carried out directly after enzymatic conversion without ssDNA purification. As in BS-seq, Cabernet measures the combined level of 5mC and 5hmC as 5mC; however, since the abundance of 5mC is much higher than 5hmC in most circumstances, the combined level of 5mC and 5hmC approximates to 5mC abundance. Additionally, omitting TET2 conversion during the first step of enzymatic treatment enables direct measurement of 5hmC, where 5hmC is specifically protected from deamination through BGT-mediated glycosylation to 5gmC and read as C while 5mC/C are deaminated and read as T, and we name this 5hmC sequencing protocol as Cabernet-H (Fig. 1a).

**Fig. 1.**
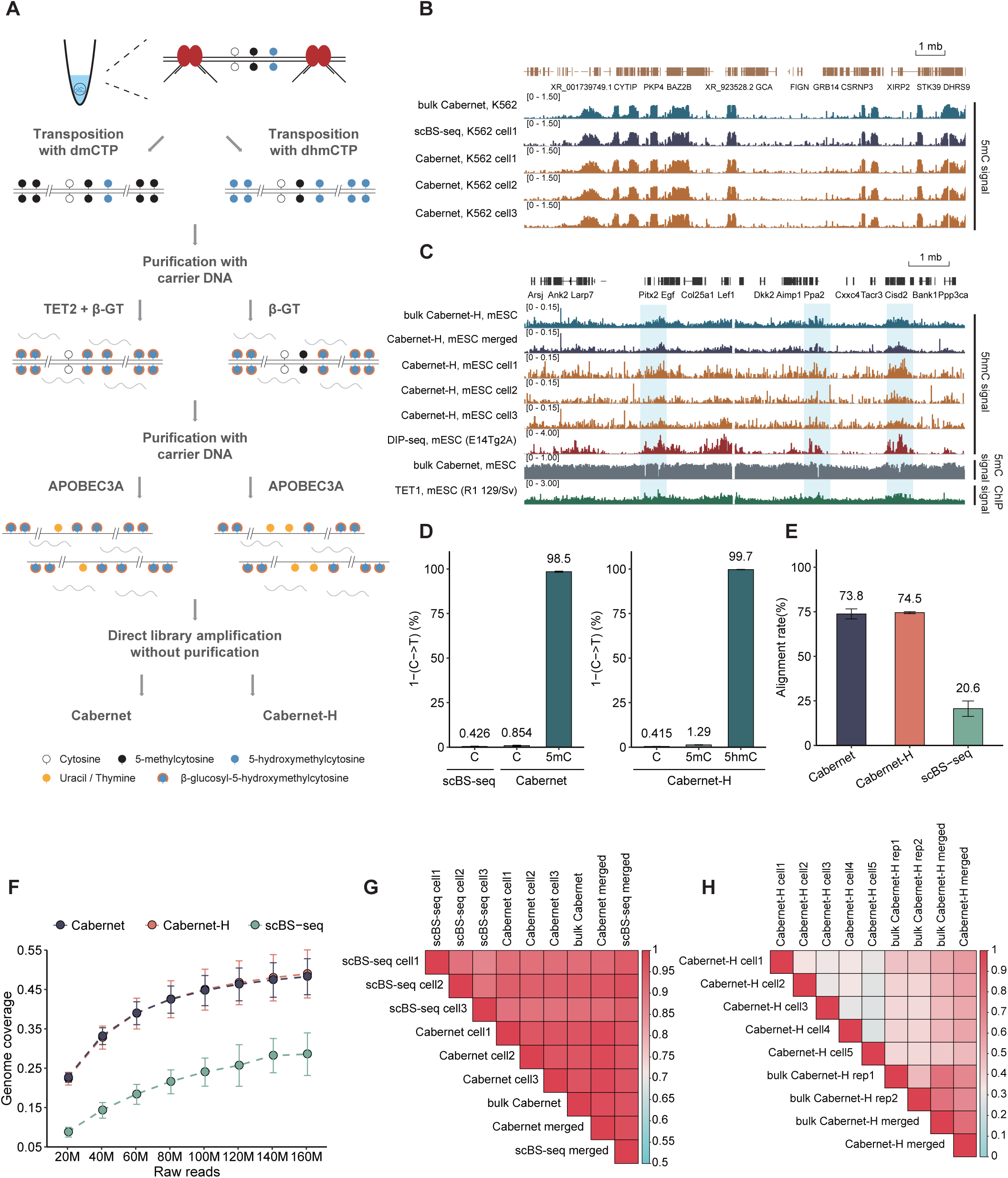
Workflow and validation of Cabernet. (*A*) Schematic of Cabernet and Cabernet-H in single cell. (*B*) IGV visualization of 5mC enrichment in K562 cell line using Cabernet sequencing method (chr2:154009136-169479494). (*C*) IGV visualization showing 5hmC enrichment in mESC using Cabernet-H sequencing method (chr3:126,231,297-137,134,000). (*D*) C-T conversion rate of 5mC in Cabernet and 5hmC in Cabernet-H; conversion rate of unmodified C is monitored by λ-DNA spike-in. (*E*) Barplot of the genome alignment rate in Cabernet, Cabernet-H and scBS-seq. (*F*) Genome coverage of Cabernet, Cabernet-H and scBS-seq under different number of downsampled reads in K562 cells. (*G*) Heatmap of Pearson correlation between Cabernet and scBS-seq sequencing results in K562. (*H*) Heatmap of Pearson correlation between single Cabernet-H and bulk Cabernet-H in mESC.

To evaluate the accuracy of Cabernet in profiling single-cell 5mC, we applied Cabernet to single K562 cells as compared with scBS-seq, which is known as the gold standard of DNA methylation detection. A high consistency was observed between Cabernet (single-cell or bulk) and scBS-seq in mapping genome-wide DNA methylation (Fig. 1b and *SI Appendix,* Fig. S1). Of note, 5mC, 5hmC, hemi-5mC or hemi-5hmC level in this study is referred to as the respective modification level at CpG sites, unless specified elsewhere. To test Cabernet-H in mapping 5hmC modifications, we applied Cabernet-H to mouse embryonic stem cells (mESCs). Genome-wide 5hmC modification map obtained by single-cell Cabernet-H and bulk Cabernet-H broadly matched previously published DIP-seq dataset (Fig. 1c and *SI Appendix,* Fig. S2) (27). As verified in K562 and mESCs, Cabernet and Cabernet-H can yield 5mC and 5hmC maps in broad match with previously proved methods.

To further validate the accuracy of Cabernet and Cabernet-H, we tested the conversion efficiency on three standard controls, which are unmodified λ-DNA, CpG methylated pUC19 plasmid, and 5hmC modified DNA fragment. In Cabernet, up to 98.5% of 5mC were correctly called, with a false-positive rate of 0.854% (C recognized as 5mC), as verified on CpG methylated pUC19 plasmid and unmodified λ-DNA. In Cabernet-H, the recall rate of 5hmC was 99.7% as verified on 5hmC modified DNA fragment, with only 0.415% of C and 1.29% of 5mC falsely called 5hmC (Fig. 1d and *SI Appendix,* Fig. S3). Besides, Cabernet and Cabernet-H also showed substantially higher mapping rates than scBS-seq (Fig. 1e), due to the use of a specific adaptor for selective amplification in contrast to the random priming strategy used by bisulfite-based methods.

The major limitation of scBS-seq is low DNA coverage, which we aim to improve with Cabernet. To examine the advantage of Cabernet over scBS-seq regarding genome coverage, we performed deep sequencing to K562 samples prepared under both protocols. The average genome coverage obtained with Cabernet and Cabernet-H was about twice that yielded by scBS-seq. Analysis of downsampled sequencing data across different depths showed Cabernet and Cabernet-H yielded a similar coverage under the same depth, both significantly outperforming scBS-seq. Specifically, Cabernet yielded near 50% genome coverage, doubling that of scBS-seq (Dataset S1). Such an advantage is especially prominent at lower sequencing depths. For example, when downsampled to 20 millions of reads (1×), the gap is amplified to 2.6 fold (Fig. 1f). Even at 0.01X∼0.1X sequencing depths, Cabernet and Cabernet-H still demonstrate an advantage (Fig. S4). This confirms that Cabernet and Cabernet-H can yield higher coverage, potentially due to less DNA damage and higher DNA recovery.

To assess the stability of Cabernet performance in single cells, we examined the correlation between single-cell measurements in homologous cell population. Pearson’s correlation analysis indicated strong correlation among single cells, revealing the stability of Cabernet. Besides, the strong correlation between single-cell Cabernet and scBS-seq further validates the accuracy of Cabernet (Fig. 1g and *SI Appendix,* Fig. S5a). By contrast, Cabernet-H analysis of mESCs (Dataset S2) demonstrated prominent 5hmC heterogeneity between individual cells, probably due to intrinsic 5hmC dynamics (Fig. 1h and *SI Appendix,* Fig. S5b). Nonetheless, merged single-cell Cabernet-H data broadly matched those obtained from bulk sequencing (Fig. 1h). As shown above, Cabernet and Cabernet-H can measure 5mC, 5hmC with high genome coverage at high accuracy.

### Cabernet-H accurately captures 5hmC

As shown above, unlike 5mC measurements, single-cell 5hmC measured by Cabernet-H does not show high consistency among single cells, probably due to intrinsic 5hmC dynamics. To further validate whether Cabernet-H faithfully captures 5hmC, we examined 5hmC distribution around the binding sites of functional proteins in mESCs. Low levels of 5mCpG and 5hmCpG were observed around the binding sites of CTCF and other methylation-averse DNA-binding proteins, such as H3K4me3 and EP300 (Fig. 2a and *SI Appendix,* Fig. S6), consistent with the DNA-binding preferences of these proteins (28, 29). In contrast, 5hmCpG was generally enriched around the center of TET1 binding sites where 5mCpG was depleted (Figs. 1c and 2b), consistent with findings that TET oxidizes 5mC to 5hmC. However, overly enriched TET proteins would lower the level of 5hmC in promoter regions (Fig. 2c and *SI Appendix,* Fig. S7a), in line with TET-mediated iterative oxidization of 5mC to 5hmC, 5fC and 5caC (30–33).

**Fig. 2.**
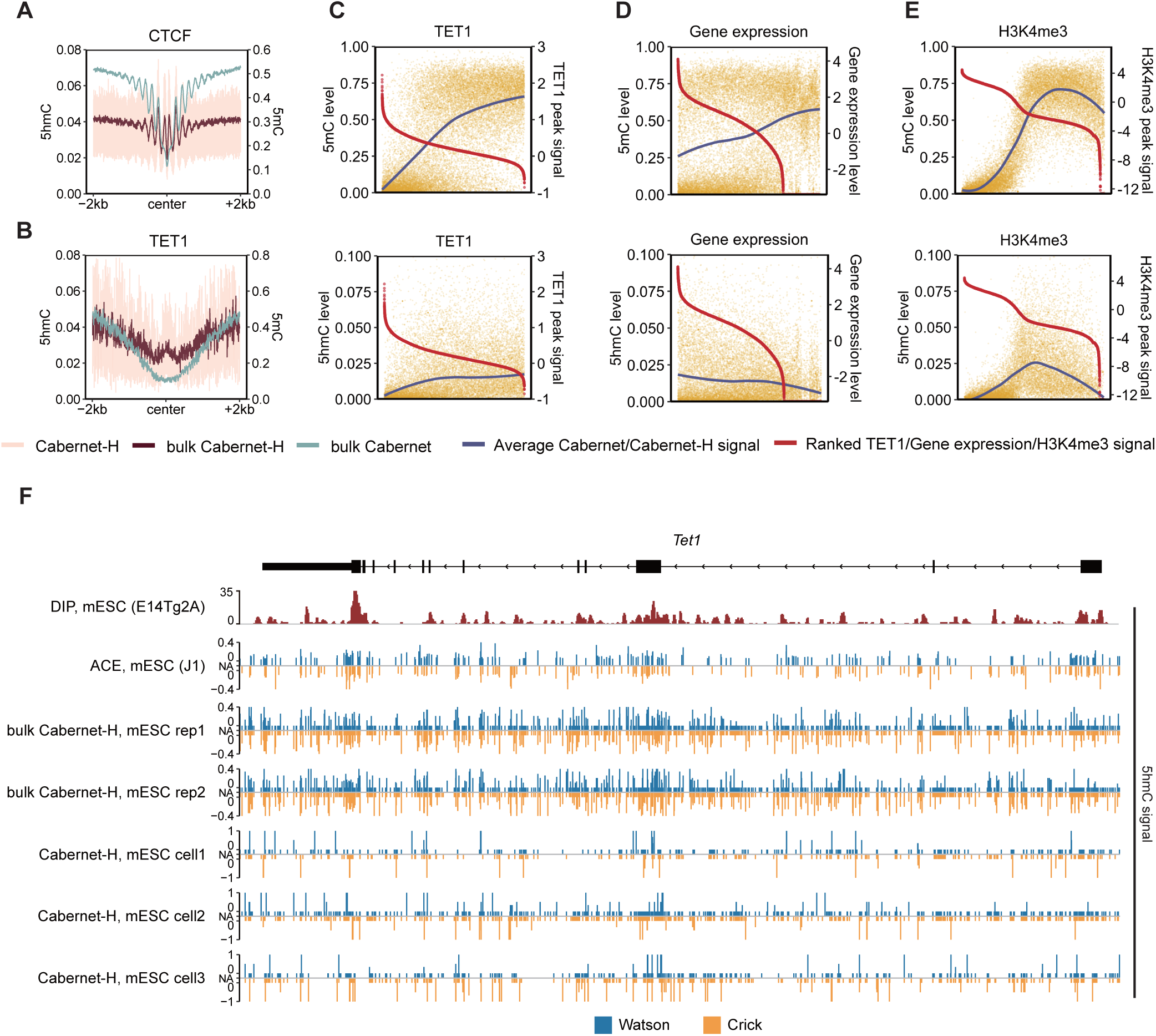
Characteristics of methylation (5mC) and hydroxy-methylation (5hmC) respectively detected by Cabernet and Caberbet-H. (*A*) Distribution of 5mC and 5hmC in the CTCF binding region (CTCF peak center ±2kb). Pink: Cabernet-H; brown: Cabernet-H_bulk; green: Cabernet_bulk. (*B*) Distribution of 5mc and 5hmC in the Tet1 binding region (CTCF peak center ±2kb). Pink: Cabernet-H; brown: Cabernet-H_bulk; green: Cabernet_bulk. (*C*) Top: signal intensities of Tet1 ChIP-Seq peaks within promoter regions in mESCs and 5mC levels at the corresponding peak regions. The red and blue fitting curves represent peak signal intensity and 5mC level in corresponding regions, respectively. Bottom: signal intensities of Tet1 ChIP-Seq peaks within promoter regions in mESCs and the 5hmC levels of the corresponding peak regions. The red and blue fitting curves represent peak signal intensity and 5hmC level in corresponding regions, respectively. The horizontal axis from left to right of each box represents the 5hmC DIP-Seq peaks, which overlapped with promoter regions, ranked by peak signal intensities from high to low. (*D*) Top: 5mC levels at promoter regions and the expression levels of corresponding genes in mESCs. The log10 of gene expression levels (transcripts per kilobase per million mapped reads, TPM) were calculated and presented. Red and blue curves represent gene expression levels and 5mC levels at promoters, respectively. Bottom: DNA 5hmC levels at promoters and the expression levels of corresponding genes in mESCs. (*E*) Top: signal intensities of H3K4me3 ChIP-Seq peaks within promoter regions in mESCs and 5mC levels at the corresponding peak regions, represented by red and blue fitting curves, respectively. Bottom: signal intensities of H3K4me3 ChIP-Seq peaks within promoter regions in mESCs and 5hmC levels at the corresponding peak regions, represented by red and blue fitting curves, respectively. (*F*) Snapshot of base-resolution 5hmC maps detected by ACE-Seq and Cabernet-H/Cabernet-H_bulk compared with DIP-Seq near the Tet1 locus (chr10:62802570-62882014; genome build: mm10). Blue and yellow represent positive (Watson) and negative (Crick) strands, respectively.

We also validated Cabernet and Cabernet-H based on the relations between 5mC/5hmC modification and gene expression. Consistent with previous reports (20, 34), our Cabernet and Cabernet-H analysis revealed that gene activity is negatively correlated with 5mC level and positively correlated with 5hmC level (Fig. 2d and *SI Appendix,* Fig. S7b). To validate the relations between 5mC/5hmC modification and histone markers, we examined 5mC and 5hmC levels in relation to H3K4me3 peak signal. 5mC depletion was observed where H3K4me3 peak signal was high. Intriguingly, 5hmC was depleted at both extreme ends of H3K4me3 peak signal intensity (Fig. 2e and *SI Appendix,* Fig. S7c). Of note, such analysis suffers a limitation that RNA-seq and ChIP-seq used a different batch of cell line than Cabernet-H.

To further validate the accuracy of 5hmC signal on a specific gene, we examined the 5hmC signal on Watson and Crick strand of Tet1 gene region captured by Cabernet-H as compared with DIP-seq and ACE-seq. It was found that 5hmC-enriched regions identified with Cabernet-H were broadly aligned with DIP-seq and ACE-seq (Fig. 2f). Besides, compared to ACE-seq, strand-asymentry was found in both bulk Cabernet-H and single-cell Cabernet-H, especially in the latter, as previously reported (24). Together, these observations reveal that Cabernet-H faithfully captures 5hmC modifications.

### 5mC and 5hmC dynamics in early mouse embryos

Having validated the feasibility and accuracy of Cabernet and Cabernet-H, we then applied them to characterize 5mC/5hmC dynamics in early mouse embryos. A total of 501 individual cells from sperms, oocytes, and pre-implantation embryos of mouse were analyzed (Dataset S3). An average of 50.4% genomic coverage was achieved for each single cell (Fig. 3a). We used Cabernet and Cabernet-H to map 5mC and 5hmC landscape across different stages during mouse embryonic development. An overall decline in 5mC level after fertilization was observed, consistent with previous findings (11, 35, 36). The level of 5hmC peaked at late zygote and late 2-cell stage, and then declined afterwards (Fig. 3b), as reported in previous studies (37–40), revealing dynamics of TET-mediated active demethylation. We subsequently merged single-cell data from each embryonic stage and analyzed the distribution of 5mC and 5hmC across the gene body, which revealed similar dynamics across developmental stages (*SI Appendix,* Fig. S8).

**Fig. 3.**
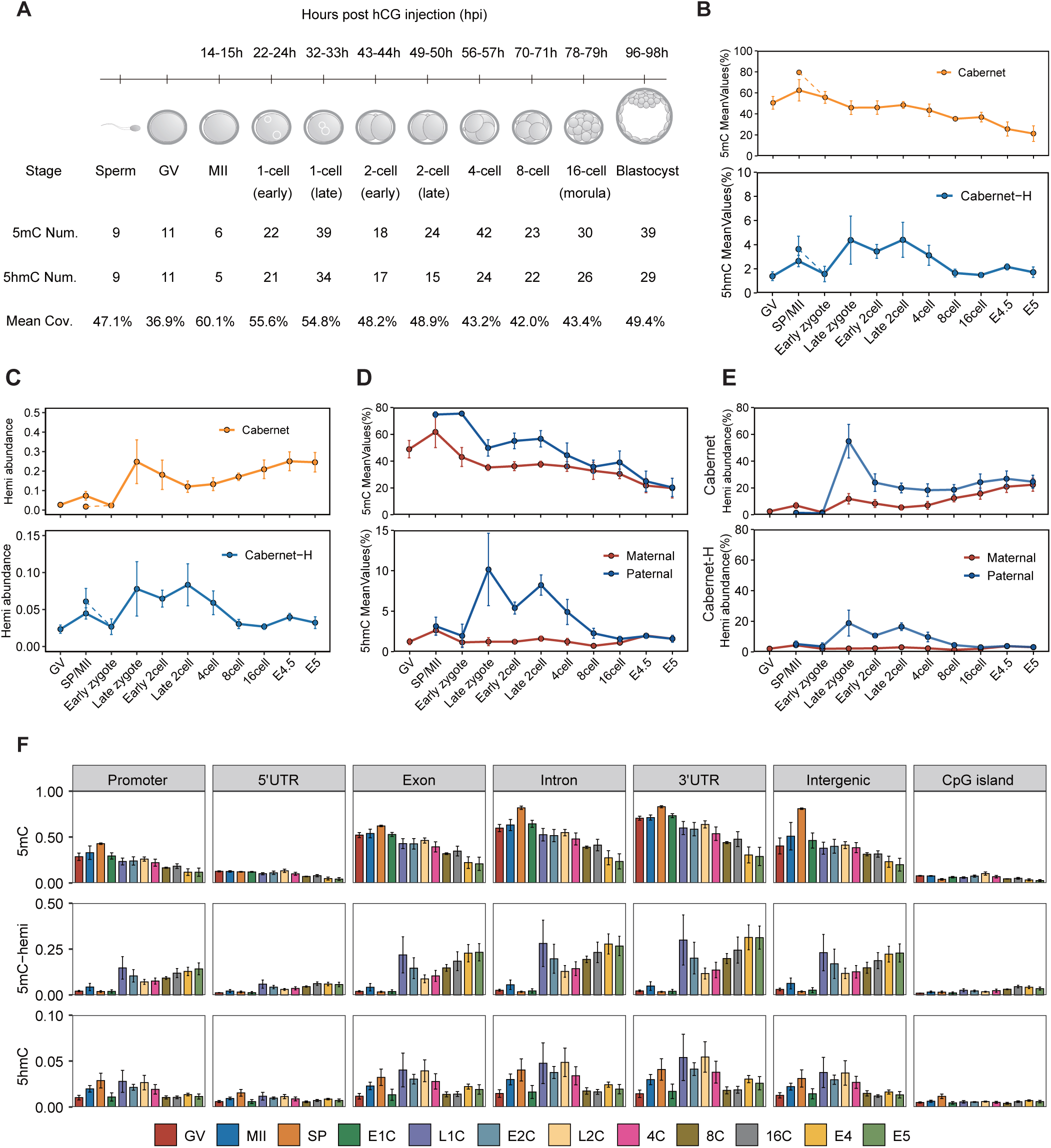
Methylation landscape across each development stage of early mouse embryos. (*A*) Experimental design. Mouse embryos at different development stages were retrieved. Each embryo was separated into single cells for further experiments with Cabernet and Cabernet-H. (*B*) Average 5mC/5hmC level at each developmental stage detected by Cabernet and Cabernet-H, respectively. (*C*) Abundance of DNA hemi-5mC (red) and hemi-5hmC (blue) across developmental stages of mouse early embryos. (*D*) Top: average 5mC level of maternal (red) and paternal (blue) genome at each developmental stage. Bottom: average DNA 5hmC level of maternal (red) and paternal (blue) genome across developmental stages of early mouse embryos. (*E*) Top: abundance of hemi-5mC in maternal and paternal genome. Bottom: abundance of hemi-5hmC in maternal (red) and paternal (blue) genome. (*F*) Top: DNA 5mC level at different genome elements in early mouse embryos. Middle: abundance of hemi-5mC within genome elements in early mouse embryos. Bottom: 5hmC level at different genomic regions across developmental stages of early mouse embryos.

Notably, since the usage of Tn5 transposome for DNA fragmentation results in different insertion sites on different alleles, the aligned reads with same start and end sites were recognized as from the same allele (*SI Appendix,* Fig. S9). Hemi-modification profiling requires measurement of the methylation status on both two DNA strands of a same allele, and only the alleles with reads sequenced from both two strands are informative of hemi-5mC and hemi-5hmC. At 1X sequencing depths, the reads of these informative alleles account for an average of 1.05% of total reads, and this ratio increases at deeper sequencing depths (Fig. S10).

In this way, hemi-5mC and hemi-5hmC landscapes in mouse embryos were mapped (Fig. 3c), which reflects passive demethylation activity (41). The abundance of hemi-5mC sharply increased after fertilization, reflecting a surge in passive demethylation; hemi-5mC abundance gradually elevated after late 2-cell when 5hmC declined (Fig. 3b,c), suggesting that passive demethylation becomes dominant after late 2-cell. Hemi-5hmC demonstrates a dynamic pattern similar to that of 5hmC, in agreement with the strand-asymmetry pattern of 5hmC distribution.

Through SNP phasing, parental differences in demethylation were able to be observed (*SI Appendix,* Fig. S11). Demethylation in the paternal genome was faster than that in the maternal genome during early embryo development, consistent with previous studies (11, 35, 42). Studies on zygote have found that 5hmC-related active demethylation is present in both maternal and paternal genome (38, 39, 43, 44). Similarly, we found that 5hmC abundance was substantially higher in paternal genome than in maternal genome, especially during late zygote and late 2-cell (Fig. 3d). To investigate if 5hmC-related active demethylation drives gene expression, we probed the 5hmC signal on the gene body of both active genes and silenced genes. It was found that for both paternal and maternal genome, 5hmC preferably enriched at the gene body of active genes instead of silenced genes (*SI Appendix,* Fig. S12), suggesting that active demethylation is a potential driver for gene activation in both paternal and maternal genome. Moreover, hemi-5mC abundance was higher in paternal genome (Fig. 3e), indicating stronger activity of passive demethylation. However, such apparently strong passive demethylation in the paternal genome appears to have little effect on gene activation, considering that hemi-5mC abundance is similar on active and silenced genes. By contrast, hemi-5mC is enriched only on active genes (*SI Appendix,* Fig. S12), suggesting that passive demethylation is associated with maternal gene activation. Together, these findings suggest different demethylation strategies for regulating paternal and maternal gene expression.

We compared 5mC, hemi-5mC and 5hmC levels at different genetic regions across different embryonic development stages. It was found that 5mC level was consistently low at 5’UTR and CpG island; genic regions (promoter, exon, intron and 3’UTR) and intergenic regions experienced notable drop in 5mC level and increase in hemi-5mC (Fig. 3f), reflecting pronounced demethylation activity after fertilization, mostly in the form of passive demethylation.

The ability of this method to measure hemi-5mC and 5hmC offers an opportunity for gaining insight into lineage differentiation during preimplantation embryonic development. Compared to 5mC, 5hmC, hemi-5mC or hemi-5hmC profile alone, integrating 5mC and hemi-5mC profiles from one single cell for t-SNE analysis would generate a more unambiguous clustering map to reveal the lineage during pre-implantation development (Fig. 4a,b,c and *SI Appendix,* Fig. S13). Unsupervised hierarchical clustering and t-SNE analysis showed that 5hmC is capable of clustering cells at different embryonic stages (Fig. 4c). Pearson’s correlation analysis revealed that batch-associated variability is much smaller than true biological variability (Fig. S14), ruling out the batch-associated confounding effects.

**Fig. 4.**
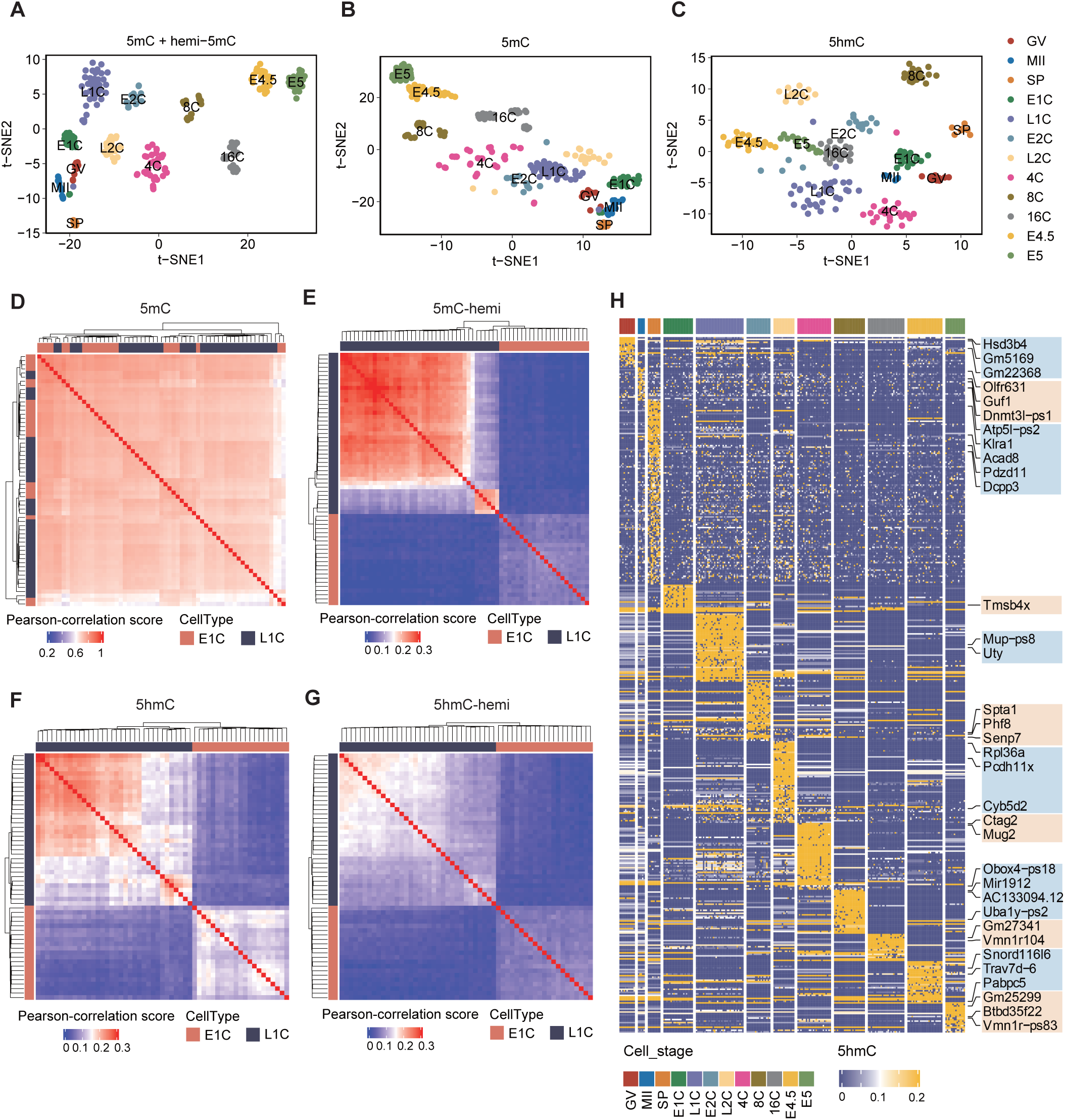
Key features of DNA methylome in early mouse embryos. (*A*) t-stochastic neighbor embedding (t-SNE) plot of cells based on the level of 5mC and hemi-5mC. Colors indicate different cell stages. (*B*) t-SNE plot of cells based on the level of 5mC. (*C*) t-SNE plot of cells based on the level of 5hmC. (*D*) Clustered heatmap showing Pearson correlation between 5mC abundance of different cells at early-1-cell stage (E1C) and late-1-cell stage (L1C). (*E*) Clustered heatmap showing Pearson correlation between hemi-5mC abundance of different cells at E1C stage and L1C stage. (*F*) Clustered heatmap showing Pearson correlation between 5hmC abundance of different cells at E1C stage and L1C stage. (*G*) Clustered heatmap showing Pearson correlation between hemi-5hmC abundance of different cells at E1C stage and L1C stage. (*H*) Abundance of 5hmC modification on promoter regions at different developmental stages.

Based on the profiles of hemi-5mC and 5hmC, different types of cells at different developmental stages can be better classified. For example, pre– and post-replication cells can be more easily distinguished from each other based on hemi-5mC profile than using 5mC profile (Fig. 4d,e and *SI Appendix,* Fig. S15a). Notably, 5hmC and hemi-5hmC profiles can also be used to differentiate pre– and post-replication cells (Fig. 4f,g and *SI Appendix,* Fig. S15b). Stage-specific 5hmC modification in promoter and gene-body regions were identified (Fig. 4h and *SI Appendix,* Fig. S15c), indicating potential regulatory functions during embryo development. Together, these suggest the capability of Cabernet and Cabernet-H for hemi-5mC, 5hmC and hemi-5hmC profiling could be used to better distinguish between cell types and states.

Here, we proved that Cabernet and Cabernet-H can be used to probe 5mC, 5hmC, hemi-5mC and hemi-5hmC at single-cell single-base resolution with high genomic coverage, providing an efficient tool for the study of biological processes during embryonic development.

### High-throughput single-cell DNA methylation sequencing with sci-Cabernet

Bisulfite treatment would limit the role of Tn5 due to DNA fragmentation, necessitating subsequent random-priming. By avoiding bisulfite treatment, Cabernet could utilize Tn5 transposome for barcoded transposition, which allows for high-throughput indexing (Fig. 5a). The technique, named sci-Cabernet, allows high-throughput single-cell methylation profiling by pooling multiple cells into one library preparation, enabling both scalability and cost-effectiveness ($1 per cell, *SI Appendix,* Fig. S16a). With this method, we were able to construct the library for thousands of single cells in 2 days by 1 person.

**Fig. 5.**
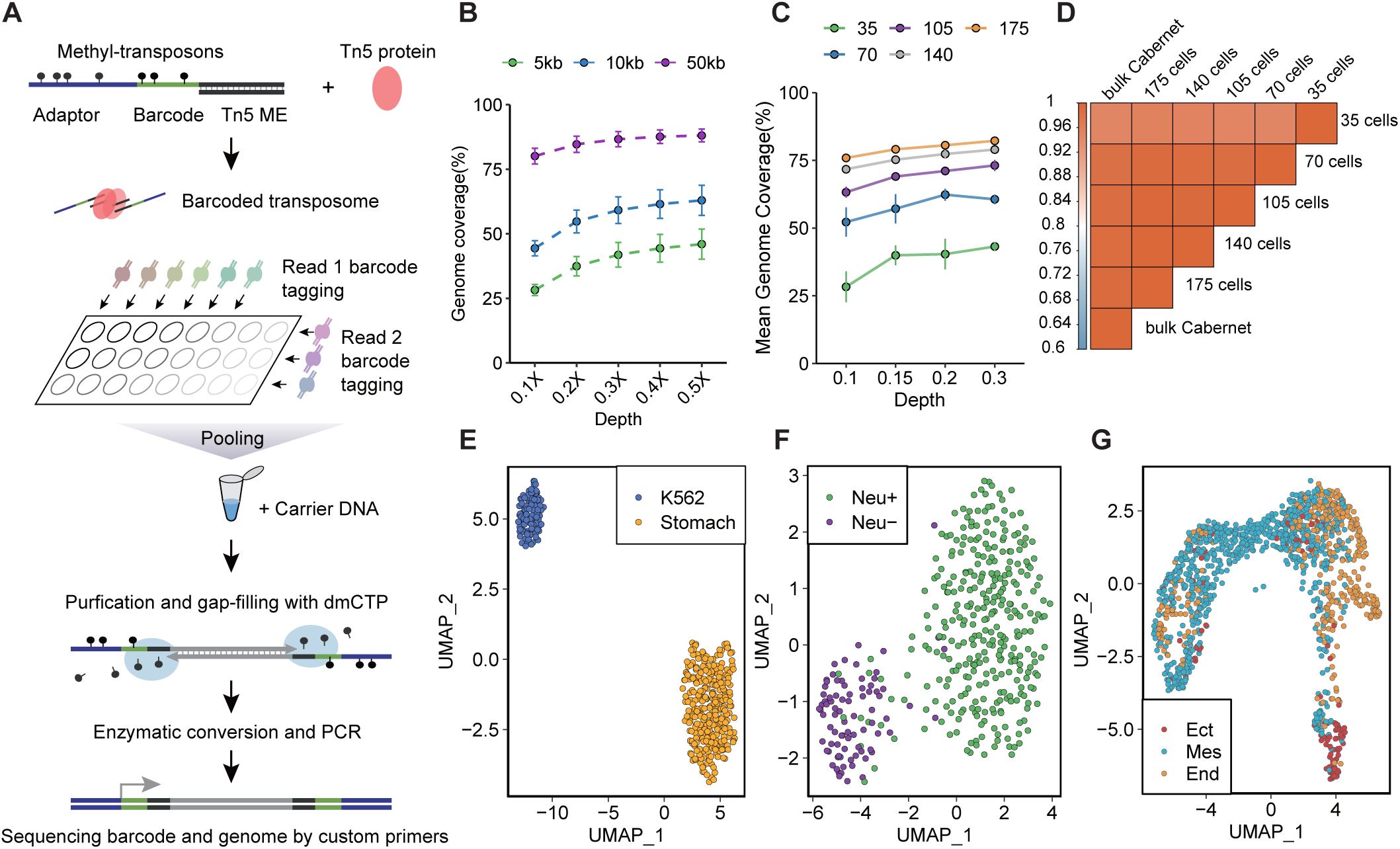
Cabernet for high-throughput single-cell DNA methylation (sci-Cabernet) profiling. (*A*) Schematic diagram of sci-Cabernet workflow. (*B*) Plot showing the genome coverage of sci-Cabernet at different sequencing depths and different bin sizes in K562 cells. (*C*) Plot showing average genome coverage of sci-Cabernet in K562 cells at different sequencing depths and different cell numbers. (*D*) Heatmap showing the correlation (Pearson correlation) of average 5mC level within 10kb using different numbers of K562 cells at 0.1X depth. (*E*) Uniform Manifold Approximation and Projection for Dimension Reduction (UMAP) figure showing the clustering of K562 cells and stomach cells after sequenced by sci-Cabernet. (*F*) UMAP showing the clustering of mouse cortical neuronal cells and non-neuronal cells after sequenced by sci-Cabernet. (*G*) UMAP visualization of E7.5 mouse embryo cells sequenced by sci-Cabernet and colored by cell-types defined by indirect annotation.

When sequenced to 0.1X average depth, sci-Cabernet could achieve 2.7 ± 0.27% genome coverage (28.2 ± 2.13% of 5-kb bin coverage), and increasing sequencing depth would lead to higher genomic coverage (Fig. 5b). We subsequently merged a few cells from the same population to perform further analysis. Merging 175 single cells can obtain about 75% genomic coverage at the 0.1× genome depth (Fig. 5c). Merged sequencing data from as few as 35 single cells broadly matched those obtained from bulk sequencing (Fig. 5d), making sci-Cabernet a reliable method for scalable single-cell DNA methylation sequencing. UMAP (Uniform Manifold Approximation and Projection) analysis suggested that two distinct cell populations as K562 cells and stomach cells could be effectively clustered after sequenced by sci-Cabernet (Fig. 5e).

We further tested the performance of sci-Cabernet on adult mouse cortex and E7.5 mouse embryo cells (Dataset S4). First, a total of 480 neurons (NeuN+) and 96 non-neuronal cells (NeuN-) from adult mouse cortex were analyzed. Based on single-cell 5mC profiles, neurons were efficiently distinguished from non-neuronal cells at high accuracy (Fig. 5f). Second, we also analyzed 5mC profiles of 1248 cells from E7.5 mouse embryo. Based on sciCabernet sequencing data, these cells were clustered into 4 groups (Fig. S16b). To determine the cell type of each group, we used FindTransferAnchors of Seuart to identify the anchors between sciCabernet sequencing data and 10× scRNA-seq data of E7.5 (GSE205117), then transferred the annotation information from scRNA-seq dataset to sciCabernet cells (Fig. S16b-c). Thus, scRNA-seq data were integrated with sciCabernet sequencing data, and scRNA-seq data were used to determine the subtypes of cells sequenced by sciCabernet. Specifically, cluster 0 is dominated by endoderm cells, cluster 1/2 is mostly comprised of mesoderm cells, and cluster 3 is largely composed of ectoderm cells. Although cell types in cluster 1 are slightly confounded, due to limitations of indirect scRNA-seq annotation, it is still obvious that mesoderm cells predominate in cluster 1 (Fig. 5g). The endoderm and mesoderm clusters are relatively adjacent to each other compared to the ectoderm cluster, which is aligned with the developmental trajectory of the three cell lineages. Together, these results demonstrate the scalability and accuracy of sci-Cabernet in cell-type determination for complex systems.

## Discussion

In this study, we have developed a carrier-assisted base-conversion by enzymatic reaction with end-tagging technology called Cabernet for base-resolution profiling of DNA methylation in single cells. With high-detectability, Cabernet stands out as a promising candidate to be broadly applied to DNA methylation analysis, especially when sample DNA quantities are extremely low such as during early embryonic development. Compared with bisulfite-based 5mC/5hmC profiling method, Cabernet uses enzymatic conversion of cytosines in the process, leading to mild reaction condition and better DNA preservation than bisulfite conversion. Compared with other bisulfite-free methods, Cabernet entails optimized purification step, minimizing the loss of target DNA in single cells whose sample DNA quantities are low in the first place. With these advantages, Cabernet can therefore capture more CpGs at a similar or even lower sequencing depth and better classify different cell types. It is also capable of mapping hemi-5mC/hemi-5hmC profile based on allele and strand assignment, allowing direct readouts of replication-dependent passive demethylation.

Although bisulfite-based single-cell 5mC mapping methods have been reported in early mouse embryos (11, 35), no previous studies have reported base-resolution mapping of 5hmC in single cells during pre-implantation development. By selectively protecting 5hmC from deamination, Cabernet-H provides an approach to effectively distinguish 5hmC from 5mC, enabling single-cell base-resolution 5hmC mapping across mouse pre-implantation development. Moreover, integrating 5mC mapping (Cabernet) with 5hmC mapping (Cabernet-H) can obtain the profiles of 5mC, 5hmC and hemi-modifications at different embryonic stages, which in turn reveals the dynamics of active and passive DNA demethylation during pre-implantation development. In addition, robust allele phasing based on parental SNP enables allele-specific 5mC and 5hmC identification. The findings suggest that TET-mediated active DNA demethylation preferentially targets paternal over maternal genome, consistent with several recently reported studies (38, 39). Lastly, we show that integrative analysis of 5mC and hemi-5mC mapping allows more precise lineage tracing across early embryonic development, and that hemi-methylation status can be used to distinguish between early and late phase of each stage.

In addition to a base-resolution mapping method for DNA methylation at single-cell level, we also provide sci-Cabernet, which employs Tn5 transposome to achieve high-throughput methylome profiling. Given the scalability and cost effectiveness of sci-Cabernet, thousands of single cell libraries can be constructed within each preparation, pointing out a reliable approach for deciphering complex biological processes such as neurogenesis and cancer development. Although the resulting genome coverage for each cell is relatively low (*SI Appendix,* Fig. S16d), merging multiple cells would substantially raise coverage (Fig. 5c). The use of enzymatic conversion instead of bisulfite treatment allows better compatability of Tn5 in Cabernet and Cabernet-H, which could enable multi-omics study that incorporates the profiling of DNA methylation and other modalities such as chromosome accessibility and 3D genome structure.

Despite its scalability, the library quality produced by sci-Cabernet may be limited by unevenness of genome coverage across different cells. The effect of high throughput was achieved by introducing a unique barcode to each individual cell via barcoded Tn5 transposomes. Tn5 transposomes with different barcodes may compete against each other during amplification, resulting in unevenness across cells. Future studies may focus on other means of introducing the barcode, such as barcode ligation.

## Materials and Methods

### Cell culture

V6.5 mESCs were cultured under feeder-free conditions in gelatin-coated plates in Dulbecco′s Modified Eagle Medium (DMEM) (GIBCO) supplemented with 15% FBS (GIBCO), 2 mM L-glutamine (GIBCO), 0.1 mM 2-mercaptoethanol (Sigma), nonessential amino acids (GIBCO), and 1,000 U/mL LIF (Millipore, ESG1107). The culture was passaged using 0.05% Trypsin (GIBCO) every 2–3 days. K562 cells were maintained in Iscove’s Modified Dulbecco’s Medium (IMDM) (Invitrogen, 12440) with 10% fetal bovine serum, 100 U/mL penicillin and 100 µg/mL streptomycin (1% P/S). Cells were split 1:8 every 2-3 days.

### Mouse embryos collection

C57BL/6N females between 6-8 weeks of age were superovulated by injection of 8-10 IU of PMSG, followed by injection of 8-10 IU of hCG 48 h later. MII oocytes were collected 14-15 h after hCG treatment. To collect early embryos, females were mated with DBA1 males after hCG injection. Early zygote was collected 14h after hCG injection and placed in medium. Different stages of embryos were harvested at 10 h (middle zygote), 16 h (late zygote), 26 h (early 2-cell), 34 h (late 2-cell), 44 h (4-cell), 51 h (8-cell), 61 h (morula) or 76 h (blastocysts) after incubation. The embryos were digested, and single cells were sorted to individual 0.2 mL Maxymum Recovery PCR tubes. Sorted single cells were then lysed in 20 mM Tris pH 8.0, 20 mM NaCl, 0.1% Triton X-100, 15 mM DTT, 1 mM EDTA, 1.5 mg/mL Qiagen protease, 0.5 μM carrier ssDNA, and 0.1 pg spike-in control at 50°C for 1 h, 65°C for 1 h, 70°C for 15 min. Lysed cells were stored at −80°C for later use.

### Transposome preparation

The transposons were designed as two versions respectively with 5mC and 5hmC modification (*SI Appendix*, Table S1). Homemade Nextera methyl-transposons have one strand of 5′-/Phos/-CTGTCTCTTATACACATCT-3′ and another strand of 5mC modified P5/P7 adaptor (Me-P5 adaptor or Me-P7 adaptor) (Genewiz, PAGE purification). Nextera hydroxymethyl-transposons have one strand of 5′-/Phos/-CTGTCTCTTATACACATCT-3′ and another strand of 5hmC modified P5/P7 adaptor (HydMe-P5 adaptor or HydMe-P7 adaptor) (Genewiz, PAGE purification). Each of the oligos was dissolved in 0.1X TE to a final concentration of 100 μM. The transposons are annealed by heating up the plates to 95°C for 1 minute and then cooling down 0.1°C per 3 seconds to reach 25°C. The transposase was purified after expressed from the pTXB1-Tn5 plasmid (Addgene). Comparable results can be generated with purchasable Tn5 transposase (Vazyme Biotech co.,ltd). Then, the transposons and transposase were dimerized into transposomes at a concentration of 1.25 μM, which was then 1:10 diluted and aliquoted for single uses and stored at −80°C.

### Preparation of spike-in control

For accurate identification of 5hmC conversion rate, cytosines in 87bp DNA fragment were modified with hydroxy-methylation by PCR amplification with 5-hydroxymethyl-dCTP supplement (ZYMO). The hydroxy-methylated DNA fragment was combined with equal amounts of CpG methylated pUC19 DNA (NEB) and unmethylated lambda DNA (NEB). CpG methylated pUC19 DNA and unmethylated lambda DNA were used to measure conversion rate of 5mC and unmodified C.

### scCabernet library preparation

Lysate from cells was transposed in a final concentration of 10 mM Tris pH 8.0, 5 mM MgCl_2_, 8% PEG 8000, 1:2,500 (0.5 nM dimer) homemade Nextera methyl-transposome and incubated at 55°C for 10 min. Transposases were removed by addition of 1 μL 2 mg/mL Qiagen protease, 1 μL 0.5 M NaCl, 75 mM EDTA and incubation at 50°C for 40 min and 70°C for 15 min. Gaps in the transposed DNA were filled by 4μL gap-filling mix (3.2 μL Q5 reaction buffer (NEB), 0.07 μL 1M MgCl2, 0.032 μL 100mM dATP, 0.032 μL 100mM dTTP, 0.032 μL 100mM dGTP, 0.32 μL 10mM 5-methyl-dCTP (NEB N0356S), 0.16 μL Q5 (NEB M0491S), 0.123 μL 120ng/μL sonicated carrier lambda DNA) and incubated at 65°C for 3 min. The product was then purified with 1.8X Ampure XP beads (Beckman Coulter). The DNA with end tags was oxidized in 15 μL TET2 reaction mix (3 μL TET2 reaction buffer plus reconstituted TET2 reaction buffer supplement (NEB), 0.3 μL oxidation supplement (NEB), 0.3 μL oxidation enhancer (NEB), 0.3 μL DTT (NEB), 1.2 μL TET2 (NEB E7125S), 1.5 μL 1:1,249 diluted Fe^2+^ solution (NEB), and 8.4 μL water) at 37°C for 1 h. Then, TET2 reaction was terminated with 0.3 μL stop solution (NEB) and incubation at 37°C for 30 min. Afterwards, the oxidized DNA was purified with 1.8X Ampure XP beads and eluted in 8 μL elution buffer (NEB). The DNA was denatured by adding 2 μL 0.1 M freshly diluted NaOH and incubation at 50°C for 10min, followed by quick cooling with ice. The APOBEC reaction mix (2 μL APOBEC reaction buffer (NEB), 0.2 μL APOBEC (NEB), 0.2 μL BSA (NEB), 1 μL 0.12 M freshly diluted HCl, and 6.6 μL water) was added to the denatured product on ice which was then incubated at 37°C for 3 h. Whole genome amplification was performed by addition of 20 μL 2X KAPA HiFi HotStart Uracil+ ReadyMix (Roche KK2801), 0.4 μL 100 μM Nextera indexed P5 primer and 0.4 μL 100 μM Nextera indexed P7 primer (*SI Appendix*, Table S2) and incubation at 98°C for 45 s, followed by 14 cycles of amplification (98°C for 15 s, 60°C for 30 s, 72°C for 1 min), and then 72°C for 3 min. Size selection was performed with SPRI beads (Beckman Coulter, typically 0.8X) according to the manufacturer’s instructions.

### scCabernet-H library preparation

Lysate was transposed in a final concentration of 10 mM Tris pH 8.0, 5 mM MgCl_2_, 8% PEG 8000, 1:2,500 (0.5 nM dimer) homemade Nextera hydroxymethyl-transposome and incubation at 55°C for 10 min. Transposases were removed by addition of 1 μL 2 mg/mL Qiagen protease, 1 μL 0.5 M NaCl, 75 mM EDTA and incubation at 50°C for 40 min, 70°C for 15 min. Transposed DNA was gap-filled by addition of 4μL gap-fill mix (3.2 μL Q5 reaction buffer (NEB), 0.07 μL 1M MgCl2, 0.08 μL 40mM dATP, 0.08 μL 40mM dTTP, 0.08 μL 40mM dGTP, 0.08 μL 40mM 5-hydroxymethyl-dCTP (ZYMO), 0.16 μL Q5 (NEB M0491S), 0.12 μL 120ng/μL sonicated carrier lambda DNA, 0.13 μL water) and incubation at 65°C for 3 min. The product was purified with 1.8× Ampure XP beads (Beckman Coulter). The end-tagged DNA was oxidized in 15 μL BGT reaction mix (0.3 μL Uridine Diphosphate Glucose (NEB), 1.5 μL NEBuffer 4 (NEB), 0.75 μL T4 Phage β-glucosyltransferase (NEB M0357S), and 12.45 μL water) at 37°C for 2 h. BGT reaction was terminated by addition of 0.75 μL Proteinase K (NEB P8107S) and incubation at 37°C for 30 min. The product was purified with 1.8× Ampure XP beads and eluted in 8 μL elution buffer (NEB). The DNA was denatured by addition of 2 μL 0.1 M freshly diluted NaOH and incubation at 50°C for 10min, followed by quick cooling with ice. The APOBEC reaction mix (2 μL APOBEC reaction buffer (NEB), 0.4 μL APOBEC (NEB), 0.2 μL BSA (NEB), 1 μL 0.12 M fresh diluted HCl, and 6.4 μL water) was added to the denatured product on ice which was then incubated at 37°C for 3 h. Whole genome amplification was performed by addition of 20 μL 2× KAPA HiFi HotStart Uracil+ ReadyMix (Roche KK2801), 0.4 μL 100 μM Nextera P5 index primer and 0.4 μL 100 μM Nextera P7 index primer and incubation at 98°C for 45 s, 14 cycles of amplification (98°C for 15 s, 60°C for 30 s, 72°C for 1 min), and 72°C for 3 min. Size selection was performed with SPRI beads (Beckman Coulter, typically 0.8 X) according to the manufacturer’s guidelines.

### Bulk Cabernet library preparation

For bulk samples, V6.5 mESCs and K562 genomic DNA was purified from 1 million cells using the QIAamp micro kit (QIAGEN), according to the manufacturer’s instructions. 3.75ng of purified genomic DNA was used for bulk Cabernet and Cabernet-H library preparation. Bulk libraries were prepared according to the protocol above, with more transposition enzyme and fewer PCR amplification cycles.

### Sci-Cabernet library preparation

Lysate in a 96-well plate was transposed in a final concentration of 10 mM Tris pH 8.0, 5 mM MgCl_2_, 8% PEG 8000, 1:3,333(0.75 nM monomer) homemade indexed P5 Nextera methyl-transposome, and 1:3,333 (0.75 nM monomer) homemade indexed P7 Nextera methyl-transposome (together forming 12×8 unique dual-index, *SI Appendix*, Table S3), and incubation at 55°C for 10 min. Transposases were removed by adding 1 μL 2 mg/mL Qiagen protease, 1 μL 0.5 M NaCl, 75 mM EDTA and incubation at 50°C for 40 min, then 70°C for 15 min. Contents in all the wells were collected and pooled together. 200ng sonicated carrier lambda DNA was added. The product was then purified with 2x 20% diluted Ampure XP beads (Beckman Coulter, diluted with PEG buffer composed of 19% PEG, 2.5M NaCl, 10mM Tris pH 8.0, 1mM EDTA, and 0.5mM Tween 20), and eluted by 20 μL 1mM Tris pH 8.0. Gaps in the transposed DNA were filled by 15μL gap-fill mix (7 μL Q5 reaction buffer (NEB), 7 μL Q5 high GC enhancer (NEB), 0.07 μL 100mM dATP, 0.07 μL 100mM dTTP, 0.07 μL 100mM dGTP, 0.7 μL 10mM 5-methyl-dCTP (NEB N0356S), 0.35 μL Q5 (NEB M0491S)) and incubation at 65°C for 3 min. The product was purified with 1.8x SPRIselect beads (Beckman Coulter), and eluted with 15 μL 1mM Tris pH 8.0. The DNA with end tags was oxidated by adding 15 μL TET2 reaction mix (6 μL TET2 reaction buffer plus reconstituted TET2 reaction buffer supplement (NEB), 0.6 μL oxidation supplement (NEB), 0.6 μL oxidation enhancer (NEB), 0.6 μL DTT (NEB), 2.4 μL TET2 (NEB E7125S), 3 μL 1:1,249 diluted Fe^2+^ solution (NEB)) at 37°C for 1 h. TET2 reaction was terminated by addition of 0.6 μL stop solution (NEB) and incubation at 37°C for 30 min. The product was purified with 1.8X SPRIselect beads and eluted in 13.5 μL 1mM Tris pH 8.0. The DNA was denatured by addition of 1.5 μL 0.2 M freshly diluted NaOH and incubation at 50°C for 10min, followed by quick cooling with ice. The APOBEC reaction mix (3 μL APOBEC reaction buffer (NEB), 0.3 μL APOBEC (NEB), 0.3 μL BSA (NEB), 1.5 μL 0.12 M fresh diluted HCl, and 9.9 μL water) was added to the denatured product on ice which was then incubated at 37°C for 3 h. Pre-amplification was performed by addition of 30 μL 2x Q5U Master Mix (NEB M0597S), 0.4 μL 100 μM pre-amplification forward primer and 0.4 μL 100 μM pre-amplification reverse primer (*SI Appendix*, Table S4) and incubation at 98°C for 30 s, 8 cycles of amplification (98°C for 10 s, 62°C for 30 s, 65°C for 90 s), and 65°C for 5 min. Excessive primers were digested with the addition of 0.5 μL Exonuclease I (NEB M0293S), and incubation at 37°C for 30 min and 72°C for 20 min. Library construction was completed by adding 15uL water, 15 μL 2x Q5U Master Mix (NEB M0597S), 0.4 μL 100 μM Nextera P5 index primer and 0.4 μL 100 μM Nextera P7 index primer, and incubation at 98°C for 30 s, 4 cycles of amplification (98°C for 10 s, 62°C for 30 s, 65°C for 90 s), and 65°C for 5 min. Purification was then done with DNA Clean & Concentrator-5 (ZYMO D4013) according to the manufacturer’s guidelines. Size selection was performed with Select-a-Size DNA Clean & Concentrator (ZYMO D4080) according to the manufacturer’s instructions.

### Deep sequencing of DNA library

PCR amplified DNA libraries were quantified by a Qubit fluorometer (Life Technologies). The libraries were pooled and quantified between the range of 150 bp and 800 bp using a Fragment Analyzer (Agilent). Library pools were sequenced on an Illumina HiSeq X Ten System, loaded at 0.9 pM, with a 10% PhiX spike-in to improve library complexity. Notably, sci-Cabernet libraries were sequenced with custom sequencing primers (*SI Appendix*, Table S5).

### Reference genome and genomic features

The ENCODE hg38 human reference genome and mm10 mouse reference genome were used in this study. Genomic blacklist regions were downloaded from ENCODE (hg38: NCFF419RSJ, mm10: ENCFF547MET). GENCODE M18 and M35 were used to annotate mouse and human genome. CGI annotation was downloaded from UCSC genome browser. Promoters are defined as regions from −2 kb to +0.2 kb relative to the transcriptional start site (TSS). The mm10 mouse genome and hg38 human genome annotated by RepeatMasker was downloaded from UCSC genome browser.

### Downloaded RNA and ChIP-seq data

A full list of the public datasets used in this study is provided in Dataset S5. The readcounts file of RNA-Seq in mESC (GSE153575) were downloaded from GEO database, and then we calculated the TPM (Transcripts Per Kilobase Million) of each gene for further analysis. The bed and bigwig file of ChIP-Seq were downloaded from ENCODE, the other ChIP-Seq data (*Mus musculus*) which were not found in ENCODE were download from GEO. First, the raw reads were processed with Trim_galore, and the cleaned data were then aligned to the reference genome; the unique mapped reads were used for further analysis. Then we used MACS2 for narrowpeak calling with default parameters. For ATAC-Seq data (GSE116854, GSE136403, GSE66581, GSE143658), after obtaining unique mapped reads, we used MACS2 to capture open chromatin regions (macs2 callpeak –t –n ––shift –75 ––extsize 150 ––nomodel –B ––SPMR –g mm ––keep-dup all).

### BS-seq sequencing data processing

Low-quality bases, short length and adaptor sequences in all single cell BS-seq and bulk BS-seq data were trimmed off using TrimGalore-0.4.5 (https://www.bioinformatics.babraham.ac.uk/projects/trim_galore/, parameters: ––fastqc ––paired –– phred33 ––retain_unpaired ––clip_R1 9 ––clip_R2 9). Cleaned reads were mapped to the hg38 human reference genome by Bismark with the following options: ‘--fastq ––non_directional –– unmapped ––nucleotide_coverage’. Reads that mapped to multiple locations were removed and PCR duplicate reads were removed with picard.jar (v-2.4.1). Methylation levels were called using Bismark methylation extractor.

### Preprocessing of Cabernet and Cabernet-H sequencing data

Low-quality bases, short length and adaptor sequences in raw sequencing reads were trimmed off using TrimGalore-0.4.5 with default parameters (https://www.bioinformatics.babraham.ac.uk/projects/trim_galore/). High-quality reads were then aligned to the reference genome (hg38 and mm10) by Bismark (v.0.19.0) (parameters: ––fastq –– non_directional ––unmapped ––nucleotide_coverage). Unmapped reads and those located in genome blacklist regions were excluded. Tn5 insertion caused 9bp gap regions then filled by methylated cytosine were removed after alignment. In this way, we were able to reduce the reads cleaning rate and improve the alignment rate. And at the same time, start and end points of the original fragments were marked. Fragments from the same allele but a different strand can be retained in the next step when removing PCR duplication reads by picard.jar (v.2.4.1). Methylation levels were called using Bismark methylation extractor.

### Preprocessing of sci-Cabernet sequencing data

Before trimming and alignment, reads of different cells should be separated from each other and barcodes and Tn5 sequences are removed in the meantime. Before separation, the barcode and Tn5 were merged to form a longer barcode sequence (referred to as barcode sequence) to improve the pick-up rate. Calculate the hamming distance between the 5’ end of the read and each barcode sequence to identify the most similar barcode indicating a specific cell. If a read has a distance of over 8 from all barcodes, then it should be filtered off. At the same time, identify the Tn5 19bp insertion site and remove the barcode sequence. Then, after the library of each single cell is established, process the single cell reads with the above pipeline described under “Preprocessing of Cabernet sequencing data”.

### 5hmC methylation calling

To call methylated sites, we counted the number of reads that supported methylation at a certain site and the number of reads that did not. These counts were then used to perform a binomial test with a probability of success equal to the conversion rate, which was determined by the fraction of methylated reads in the lambda genome (spike-in during library construction). The false discovery rate (FDR) for a given P-value cutoff was computed using Benjamini–Hochberg approach.

### Calling of hemi-5mC/hemi-5hmC abundance at CpGs

The use of Tn5 transposome for DNA fragmentation would result in different insertion sites on different alleles. If two reads are aligned to the same starting and ending sites on the reference genome, then they are regarded as from a same allele. First, select those alleles that have reads sequenced from both the Watson strand and the Crick strand. Then, measure the methylation status (5mC/5hmC) at CpG dinucleotides on both strands of the same allele, and calculate the number of hemi-methylated CpGs (hemi-5mCpGs, CpGs wherein one strand has a methylated C and the other does not) and hemi-hydroxymethylated CpGs (hemi-5hmCpGs, CpGs wherein one strand has a hydroxymethylated C and the other does not). Hemi-5mC/hemi-5hmC abundance was calculated as the ratio of hemi-5mCpGs/ hemi-5hmCpGs to the total number of CpGs detected on two strands.

### Parental phasing

The reference SNPs of C57BL/6N and DBA/1J were downloaded from Mouse Genomes Project (REL-1505-SNPs_Indels) which can be accessed on https://www.sanger.ac.uk/data/mouse-genomes-project/. Only the SNPs that could distinguish the paternal (DBA/1J) and maternal (C57BL/6N) genomes were used in our analysis. Considering that the C->T transition cannot distinguish the allele of C and T, so in the case of C->T transition, remove the C and T appearing on the same allele. The rest of SNPs in the genome were used to classify the reads into paternal and maternal reads. The same also applies to G->A transition. Only those SNPs with base quality >30 are included in the analysis. When the difference value (D-Values) between the ratio of SNPs on a read belonging to paternal SNPs and the ratio belonging to maternal SNPs is over 0.2, designate the reads as maternal or paternal. The aligned reads were further processed to calculate the methylation levels at CpG sites.

### Correlation between Cabernet and scBS-seq datasets

The average methylation levels within 10kb windows were calculated by deepTools(45) multiBigwigSummary. To profile the relationship between the results from scBS-seq and Cabernet method, Pearson correlation coefficient (PCC) was calculated directly from the average 5mC/5hmC level within 10kb windows of K562 cell line detected by scCabernet, bulk Cabernet, merged scCabernet, scBS-seq and scBS-seq merged datasets. At the same time, PCC was also calculated between scCabernet-H, bulk Cabernet-H and merged scCabernet-H with mESC cells.

### Methylation distribution across gene body and ChIP-peak center

The average methylation levels around genic regions (from 2kb upstream of TSS through the gene body to 0.5kb downstream of the transcription end sites) were computed for each gene in each dataset by computeMatrix scale-regions. Similarly, the average methylation levels around ChIP-peak region, which is between 2kb upstream and downstream of the peak center, were computed by computeMatrix reference-point. Values obtained from cells of the same stage or cell types were combined to plot the average levels around the genomic and ChIP-peak regions.

### Identifying highly methylated regions and dimension reduction clustering

The average 5mC/5hmC levels within promoter regions (–2kb, TSS, +0.2kb) and gene body regions (from TSS to TES) were computed for each dataset by computeMatrix BED-file as methylation features. Seurat packages with FindAllMarkers function were used to find the specific highly methylated regions during each stage. The heat map of these highly methylated regions was drawn by ComplexHeatmap (46). The top 8000 features calculated by Seurat FindVariableFeatures function were used to perform dimension reduction clustering by Rtsne (47).

### Correlation between cell stages

Pearson correlation coefficient (PCC) was calculated between cells of E1C and L1C based on methylation levels on promoter regions. First, calculate the average methylation (5mC/5hmC/hemi-5mC/hemi-5hmC) levels at promoter regions (–2kb, TSS, +0.2kb) for each cell of zygote stage (E1C and L1C). Then calculate PCC between each two different cells and draw the heat map using ComplexHeatmap.

## Acknowledgments

This study is supported by Beijing Advanced Innovation Center for Genomics at Peking University and the Beijing Municipal Commission of Science and Technology (Z201100005320015). We thank the staff at High-throughput Sequencing Center, Peking University for their help in next-generation sequencing. We thank all the group members for their insightful discussion on this project.

---

**Fig. S1.**
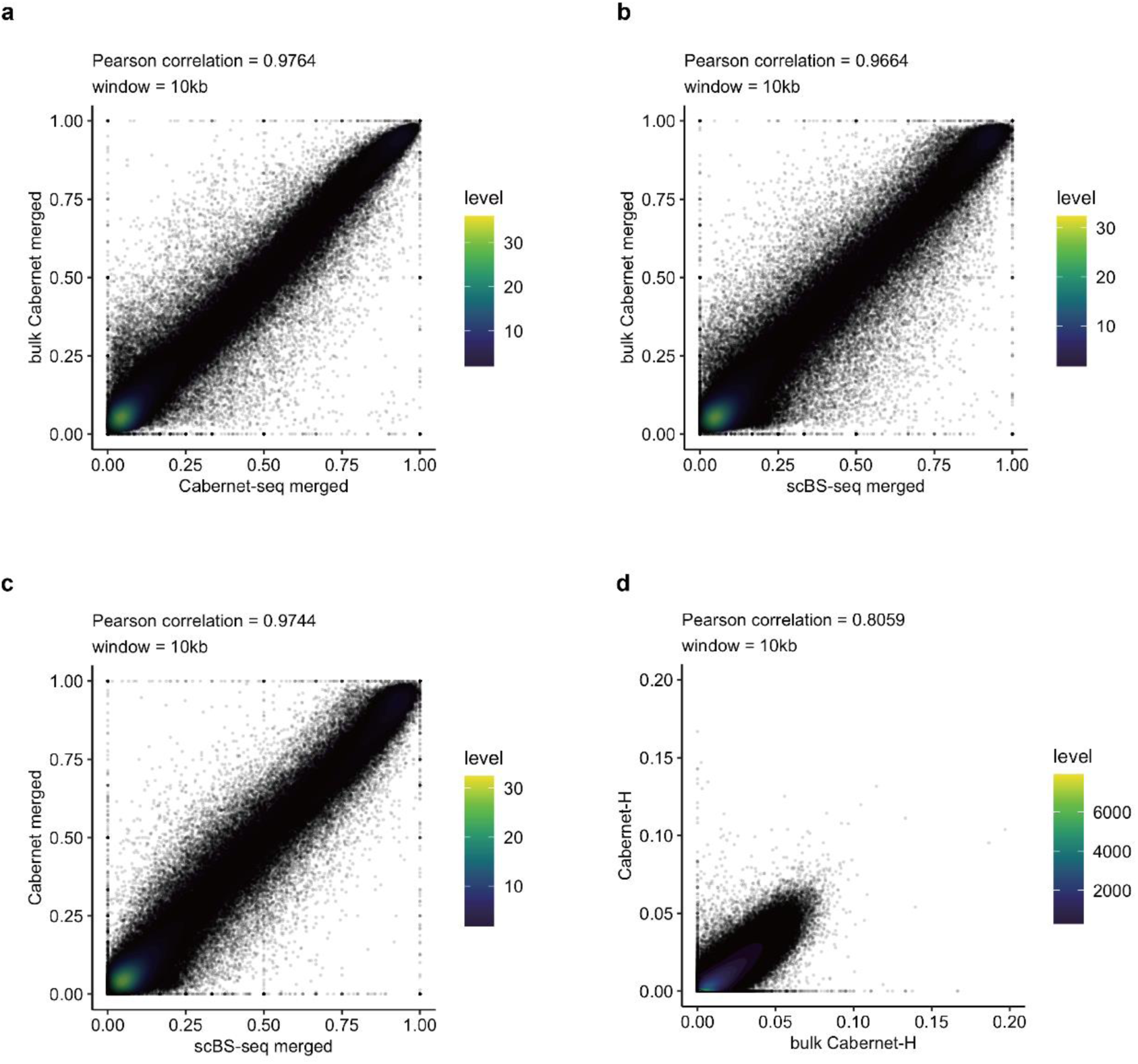
Correlation between different methylome sequencing methods. (**a**) Correlation density plot of signals between Cabernet (merged from single cells) and Cabernet_bulk (bulk) in K562 cells. Correlation analysis was performed with 10-kb bins spanning the genome. **(b)** Correlation density plot between Cabernet_bulk signals and scBS-seq signals in K562 cells (in 10-kb bins). **(c)** Correlation density plot between Cabernet signals and scBS-seq signals in K562 cells (in 10-kb bins). **(d)** Correlation density plot of Cabernet-H signals (single cell) and Cabernet-H_bulk (bulk) in mESCs (in 10-kb bins).

**Fig. S2.**
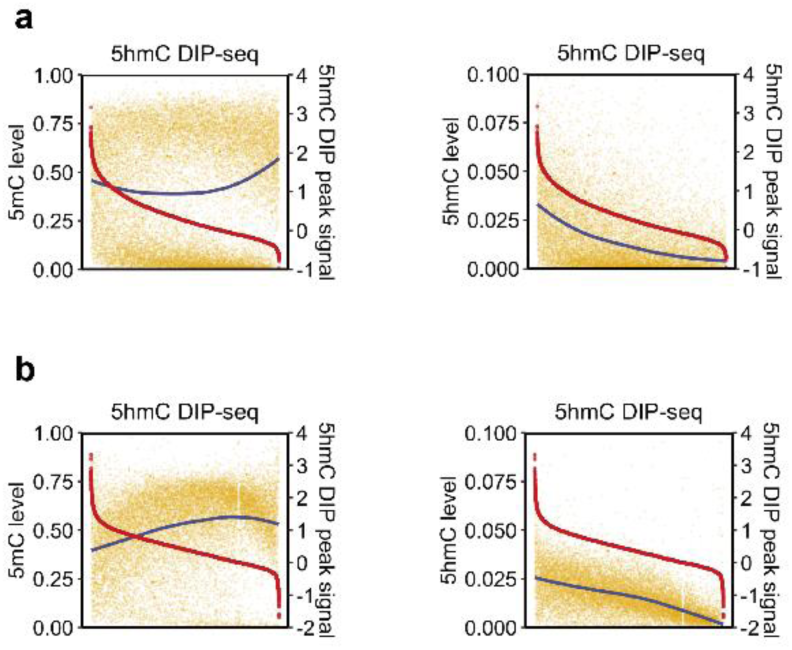
(**a**) Left: signal intensities of 5hmC DIP-Seq peaks (red) within promoter regions in mESCs and 5mC levels detected by Cabernet (blue) at the corresponding peak regions. Right: signal intensities of 5hmC DIP-Seq peaks (red) within promoter regions in mESCs and 5hmC levels (blue) at the corresponding peak regions. **(b)** Left: signal intensities of 5hmC DIP-Seq peaks (red) within gene body regions in mESCs and 5mC levels detected by Cabernet (blue) at the corresponding peak regions. Right: signal intensities of 5hmC DIP-Seq peaks (red) within gene body regions in mESCs and 5hmC levels (blue) at the corresponding peak regions.

**Fig. S3.**
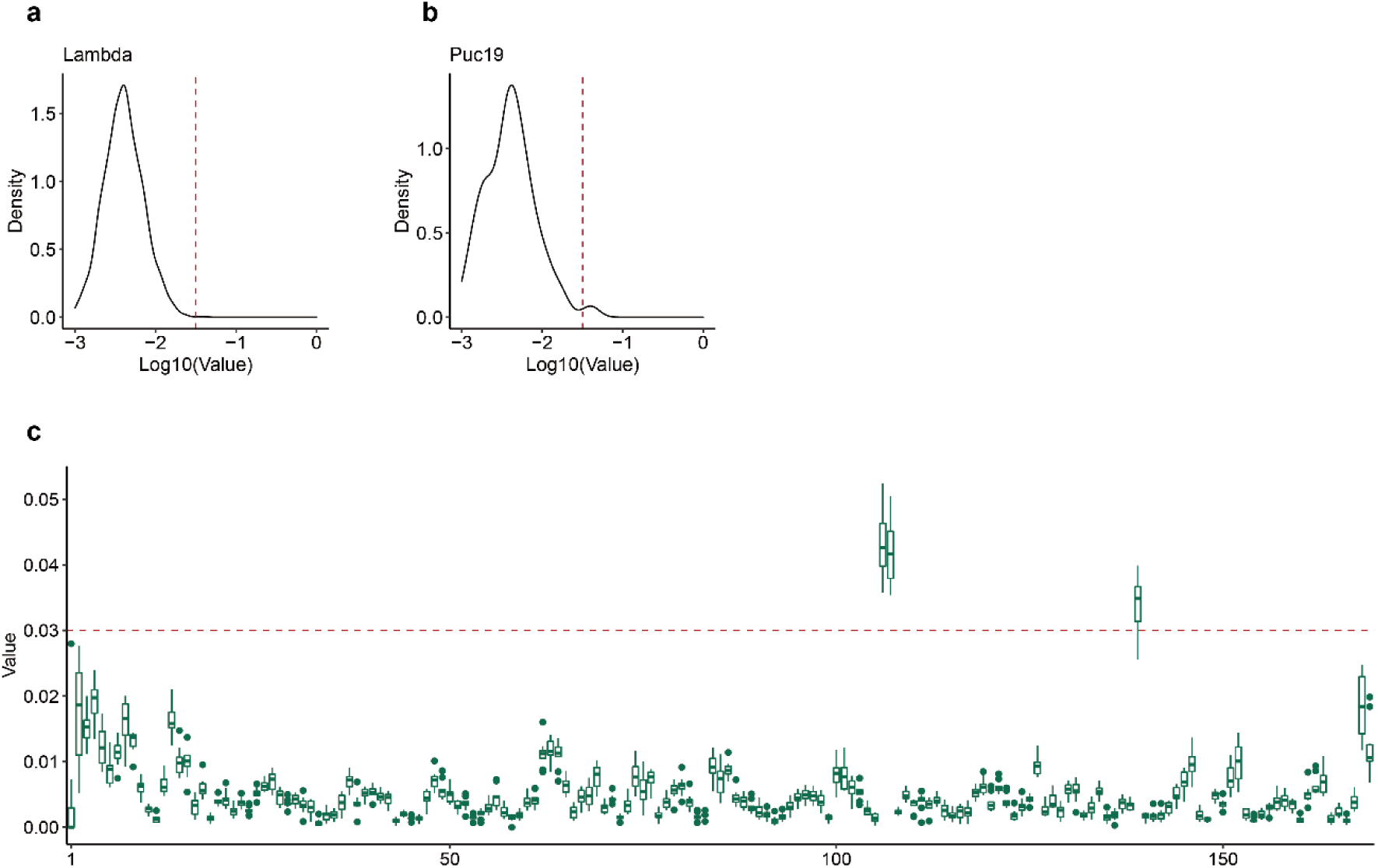
Overview of Cabernet-H data quality. (**a**) Density distribution of 5hmC meanvalue detected by Cabernet-H in lambda DNA. The read dotted line is at log10(0.03). **(b)** Density distribution of 5hmC meanvalue detected by Cabernet-H in puc19. The read dotted line is at log10(0.03). **(c)** Meanvalue of 5hmC at each CpG sites detected by Cabernet-H in puc19.

**Fig. S4.**
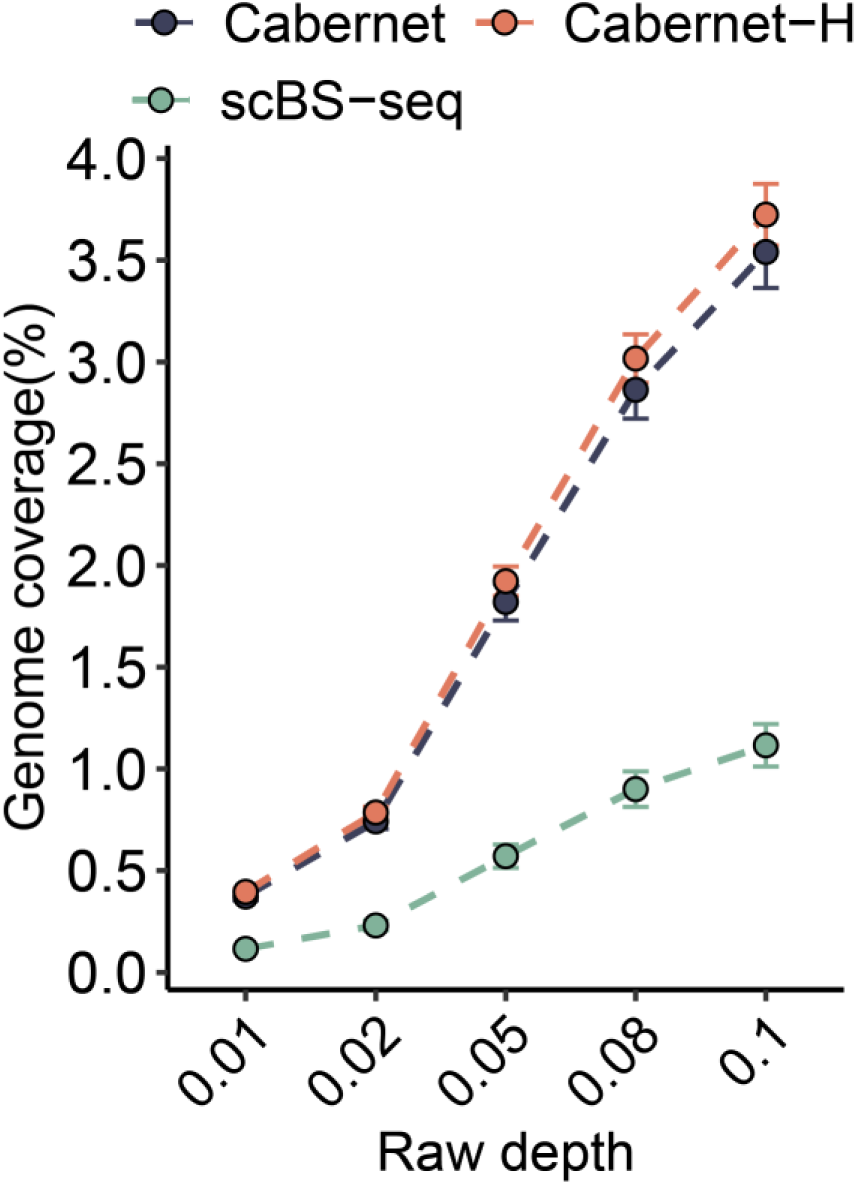
Genome coverage of Cabernet, Cabernet-H and scBS-seq under different number of downsampled reads in K562 cells.

**Fig. S5.**
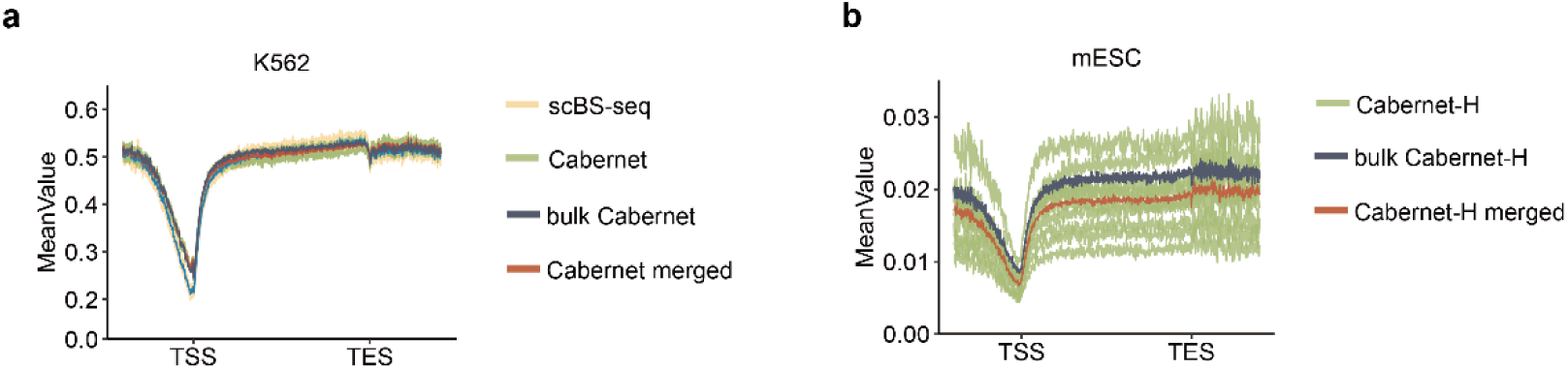
Modification pattern of 5mC and 5hmC detected by Cabernet and Cabernet-H on gene body. (**a**) Modification pattern of 5mC detected by Cabernet and scBS-seq on gene body in K562 cells. Averaged DNA methylation levels along gene bodies between 2 kilobase (kb) upstream of the transcription start sites (TSS) and 2 kb downstream of the transcription end sites (TES) of all RefSeq genes. Green: Cabernet; red: Cabernet_merged; gray: Cabernet_bulk; yellow: scBS-seq. **(b)** Modification pattern of 5hmC detected by Cabernet-H on gene body in mESCs. Averaged 5hmC levels along the gene bodies between 2 kb upstream of the transcription start sites (TSS) and 2 kb downstream of the transcription end sites (TES) of all RefSeq genes. Green: Cabernet-H; red: Cabernet-H_merged; gray: Cabernet-H_bulk.

**Fig. S6.**
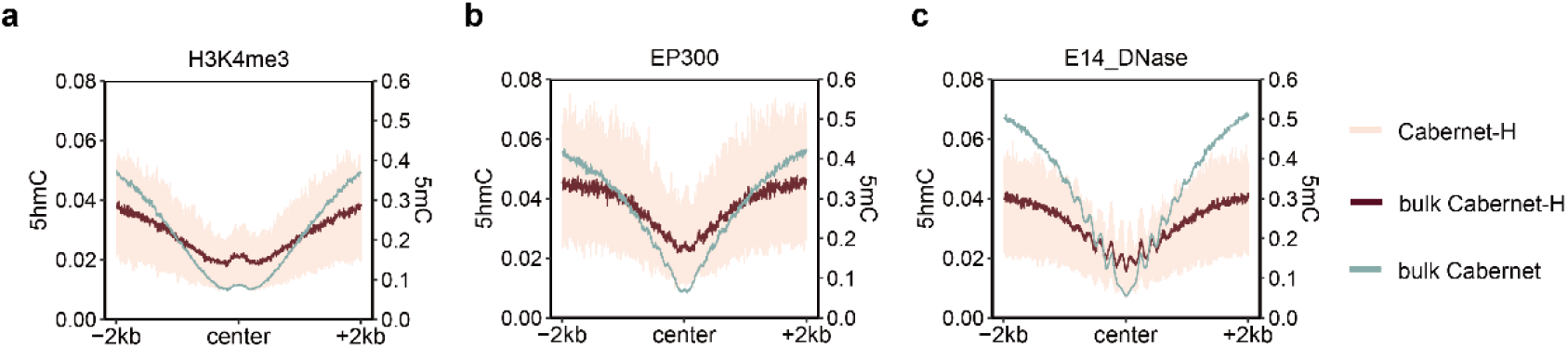
Enrichment results of Cabernet and Cabernet-H at binding regions of different factors. (**a**) Enrichment characteristics of Cabernet and Cabernet-H at H3K4me3 binding region (H3K4me3 peak center ±2kb). Pink: Cabernet-H; brown: Cabernet-H_bulk; green: Cabernet_bulk. **(b)** Enrichment characteristics of Cabernet and Cabernet-H at EP300 binding region (EP300 peak center ±2kb). Pink: Cabernet-H; brown: Cabernet-H_bulk; green: Cabernet_bulk. **(c)** Enrichment characteristics of Cabernet and Cabernet-H at E14_DNase binding region (DNase peak center ±2kb).

**Fig. S7.**
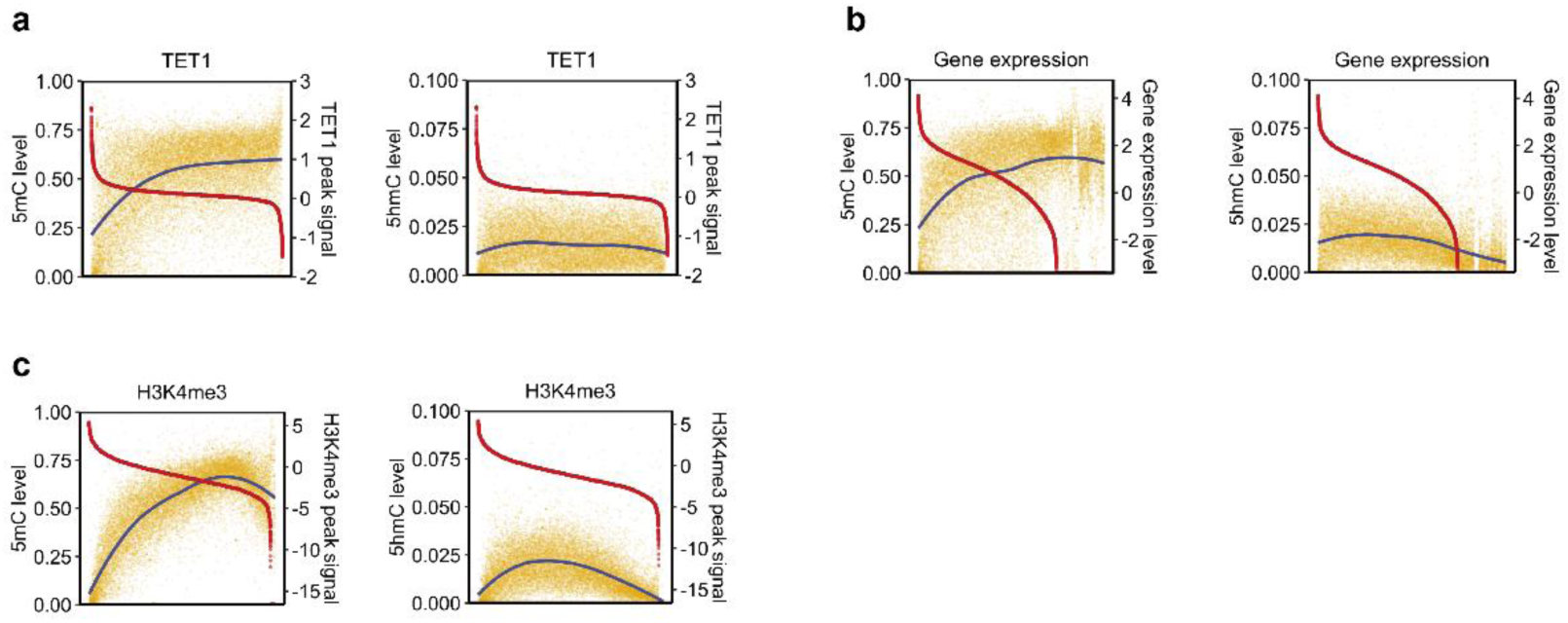
Comparison between 5mC/5hmC levels detected by Cabernet/Cabernet-H and signal intensities of different factors at gene body region. (**a**) Left: signal intensities of Tet1 ChIP-Seq peaks (red) within gene body regions in mESCs and the 5mC levels detected by Cabernet (blue) at the corresponding peak regions. The horizontal axis from left to right of each box represents the Tet1 peaks, which overlapped with gene body regions, ranked by peak signal intensities from high to low. Right: signal intensities of Tet1 ChIP-Seq peaks within gene body regions in mESCs and the DNA 5hmC levels at the corresponding peak regions. **(b)** Left: 5mC levels (blue) at gene body regions and the expression levels of corresponding genes (red) in mESCs. The log10 of gene expression levels (transcripts per kilobase per million mapped reads, TPM) were calculated and presented. Right: DNA 5hmC levels at gene body regions and the expression levels of corresponding genes in mESCs. **(c)** Left: signal intensities of H3K4me3 ChIP-Seq peaks (red) within gene body regions in mESCs and 5mC levels (blue) at the corresponding peak regions. Right: signal intensities of H3K4me3 ChIP-Seq peaks (red) within gene body regions in mESCs and 5hmC levels (blue) at the corresponding peak regions.

**Fig. S8.**
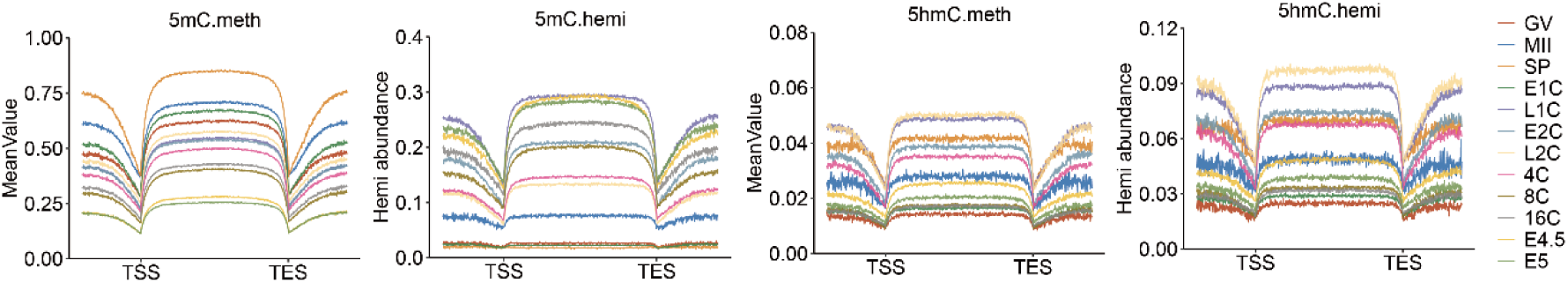
Key features of DNA methylome in early mouse embryos. Distribution of 5mC/hemi-5mC, 5hmC/hemi-5hmC detected by Cabernet/Cabernet-H in early mouse embryos on gene body regions between 2 kilobase (kb) upstream of the transcription start sites (TSS) and 2-kb downstream of the transcription end sites (TES) of all RefSeq genes. Different colors represent different stages during embryonic development.

**Fig. S9.**
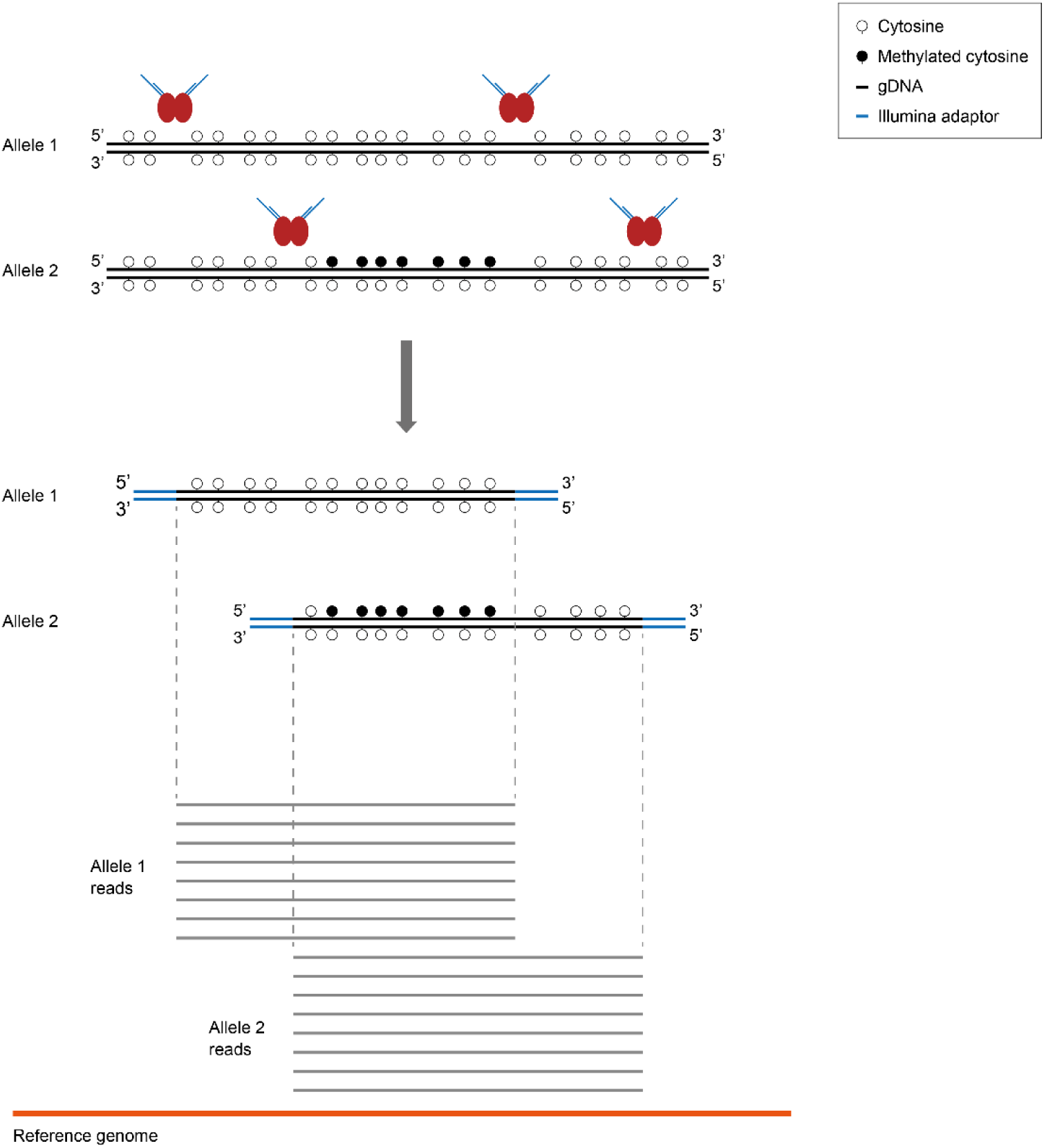
Principle for allele counting in Cabernet. The amplicons aligned to the same starting and ending sites on the reference genome are originated from the same allele of the single-cell genomic DNA. This allows for the detection of hemi-methylation in each allele.

**Fig. S10.**
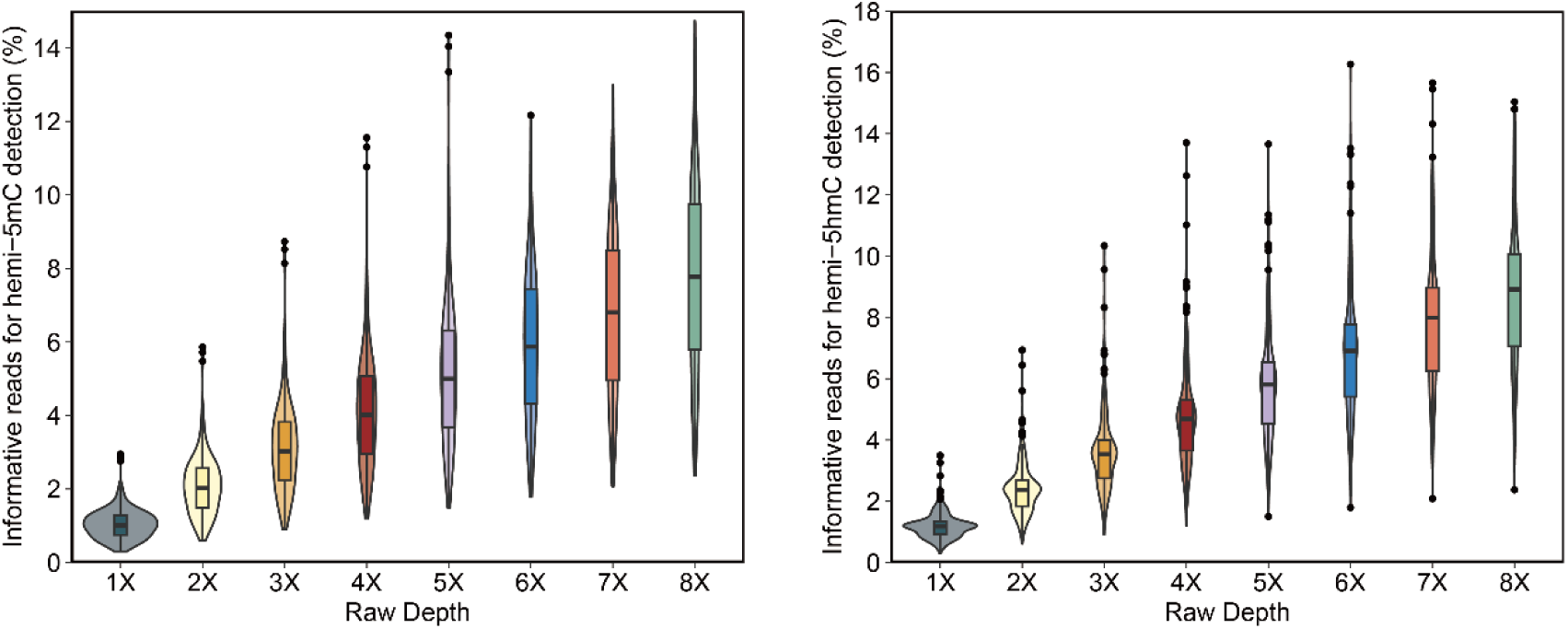
Average fraction of reads from each cell that are informative of hemi-5mC and hemi-5hmC at varied sequencing depths.

**Fig. S11.**
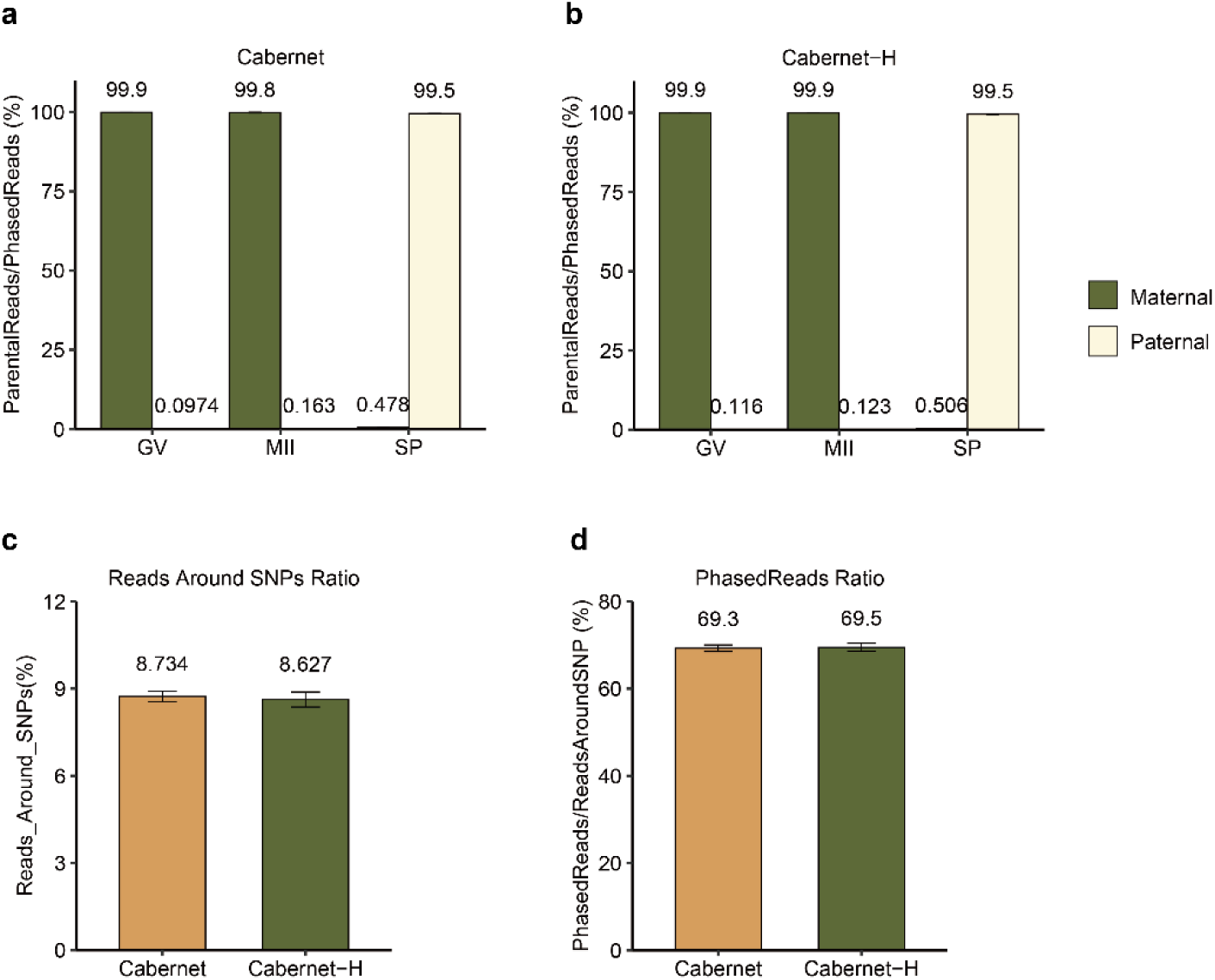
Accuracy of parental phasing for Cabernet and Cabernet-H. (**a**) The ratio of maternal/ paternal reads to all phased reads in GV/MII/SP cells sequenced by Cabernet. **(b)** The ratio of maternal/ paternal reads to all phased reads in GV/MII/SP cells sequenced by Cabernet-H. **(c)** The percentage of reads around SNPs in all reads. **(d)** The ratio of phased reads to all reads around SNPs.

**Fig. S12.**
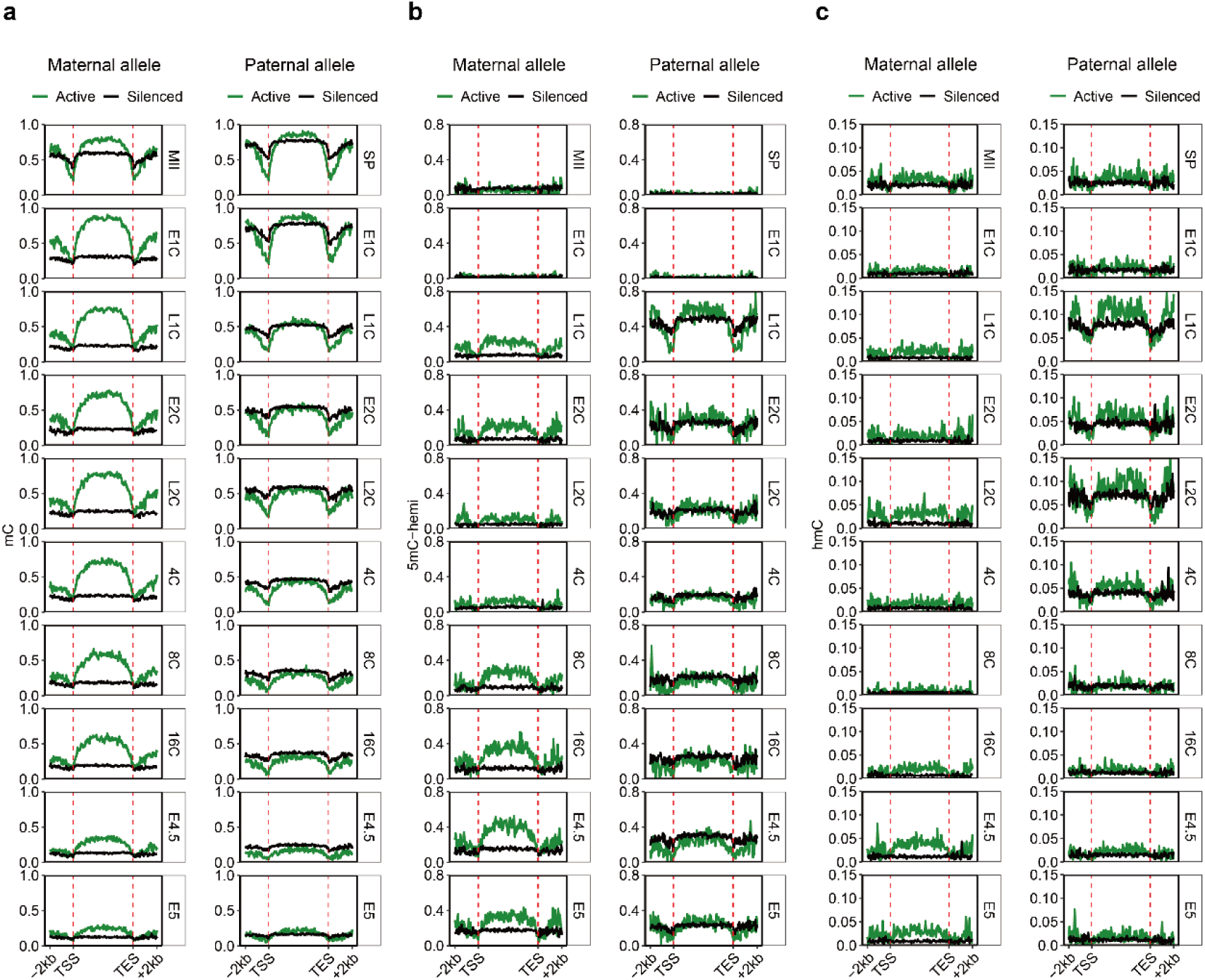
Methylation levels at gene body of active/silenced genes in the maternal/paternal genome of oocytes/sperm cells and early mouse embryos. (**a**) 5mC level at gene body of active/silenced genes in the maternal (left) and paternal genome (right). **(b)** hemi-5mC abundance at gene body of active/silenced genes in the maternal (left) and paternal genome (right). **(c)** 5hmC level at gene body of active/silenced genes in the maternal (left) and paternal genome (right).

**Fig. S13.**
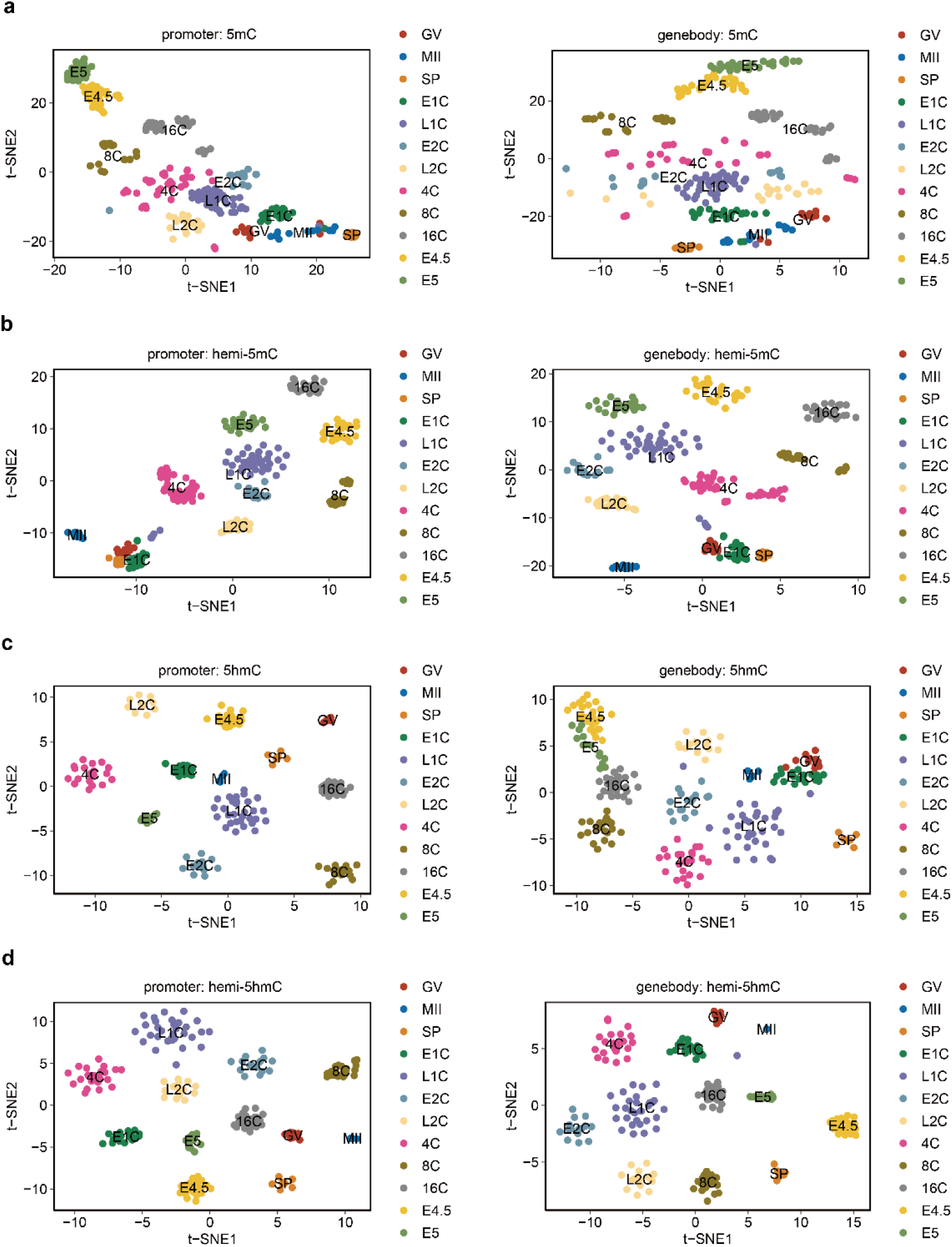
t-SNE analysis of cells based on 5mC/5hmC/hemi-5mC/hemi-5hmC. Colors indicate different cell stages. (**a**) t-SNE plot of 5mC level at promoter (left) and gene body (right). **(b)** t-SNE plot of hemi-5mC abundance at promoter (left) and gene body (right). **(c)** t-SNE plot of 5hmC level at promoter (left) and gene body (right). **(d)** t-SNE plot of hemi-5hmC abundance at promoter (left) and gene body (right).

**Fig. S14.**
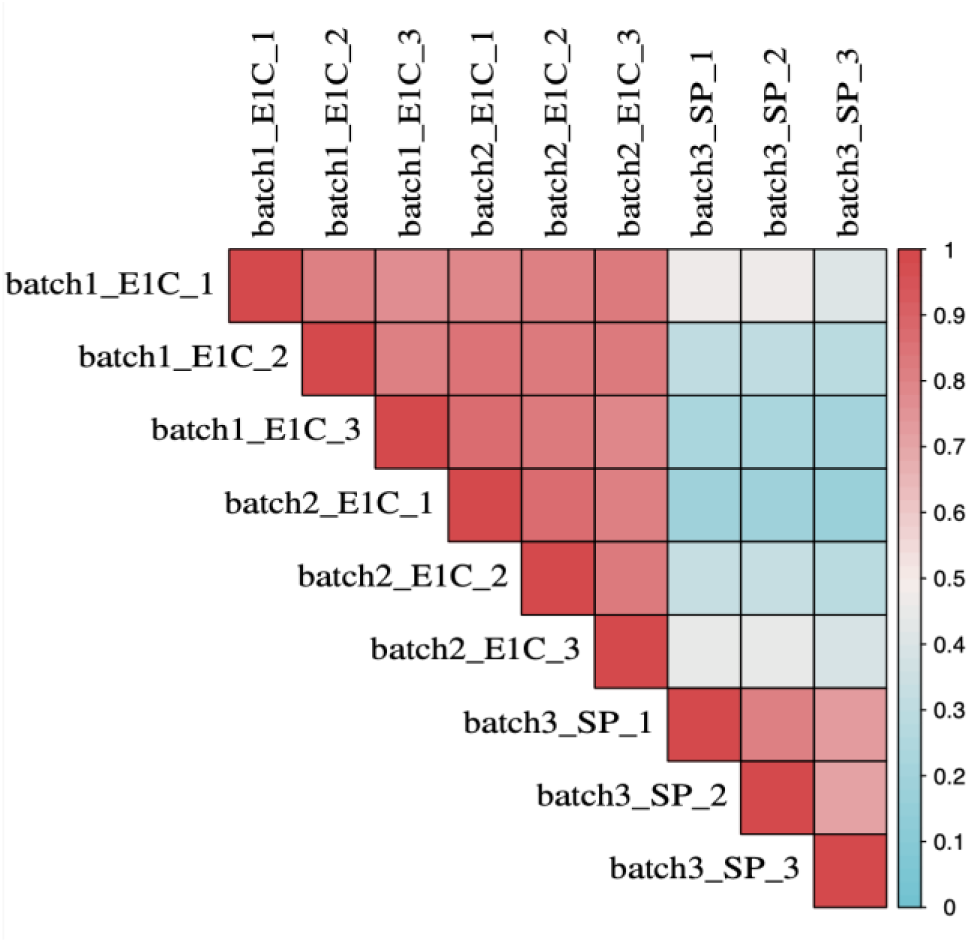
Heatmap of Pearson correlation of 5mC levels between two different batches of early 1-cells and sperm cells.

**Fig. S15.**
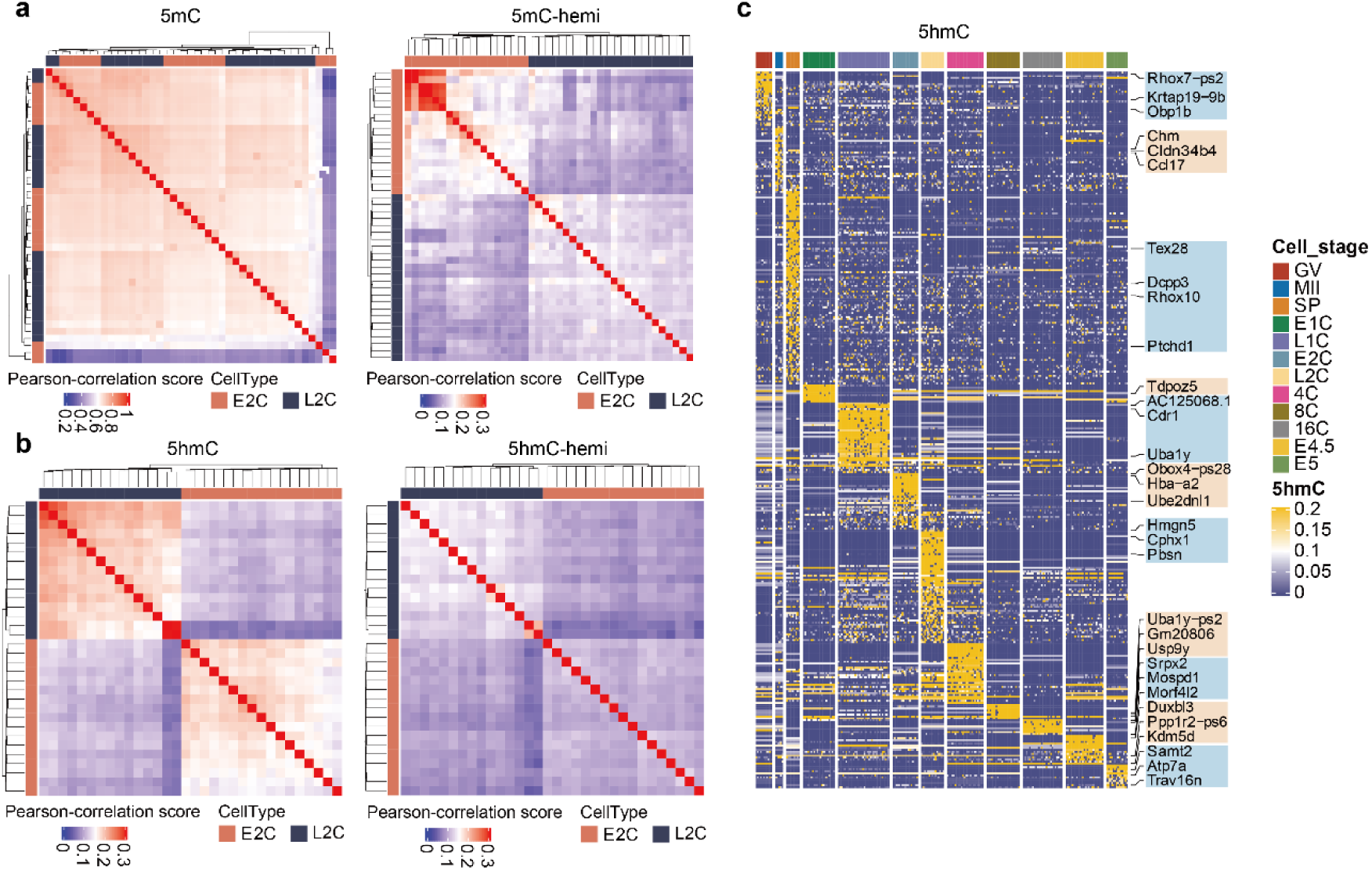
Key features of DNA methylome during early mouse embryos. (**a**) Left: clustered heatmap showing Pearson correlation between 5mC abundance of different cells at early-2-cell stage (E2C) and late-2-cell stage (L2C). Right: clustered heatmap showing Pearson correlation between hemi-5mC abundance of different cells at E2C stage and L2C stage. **(b)** Left: clustered heatmap showing Pearson correlation between 5hmC abundance of different cells at E2C stage and L2C stage. Right: clustered heatmap showing Pearson correlation between hemi-5hmC abundance of different cells at E2C stage and L2C stage. **(c)** Abundance of 5hmC modification on gene body regions at different developmental stages.

**Fig. S16.**
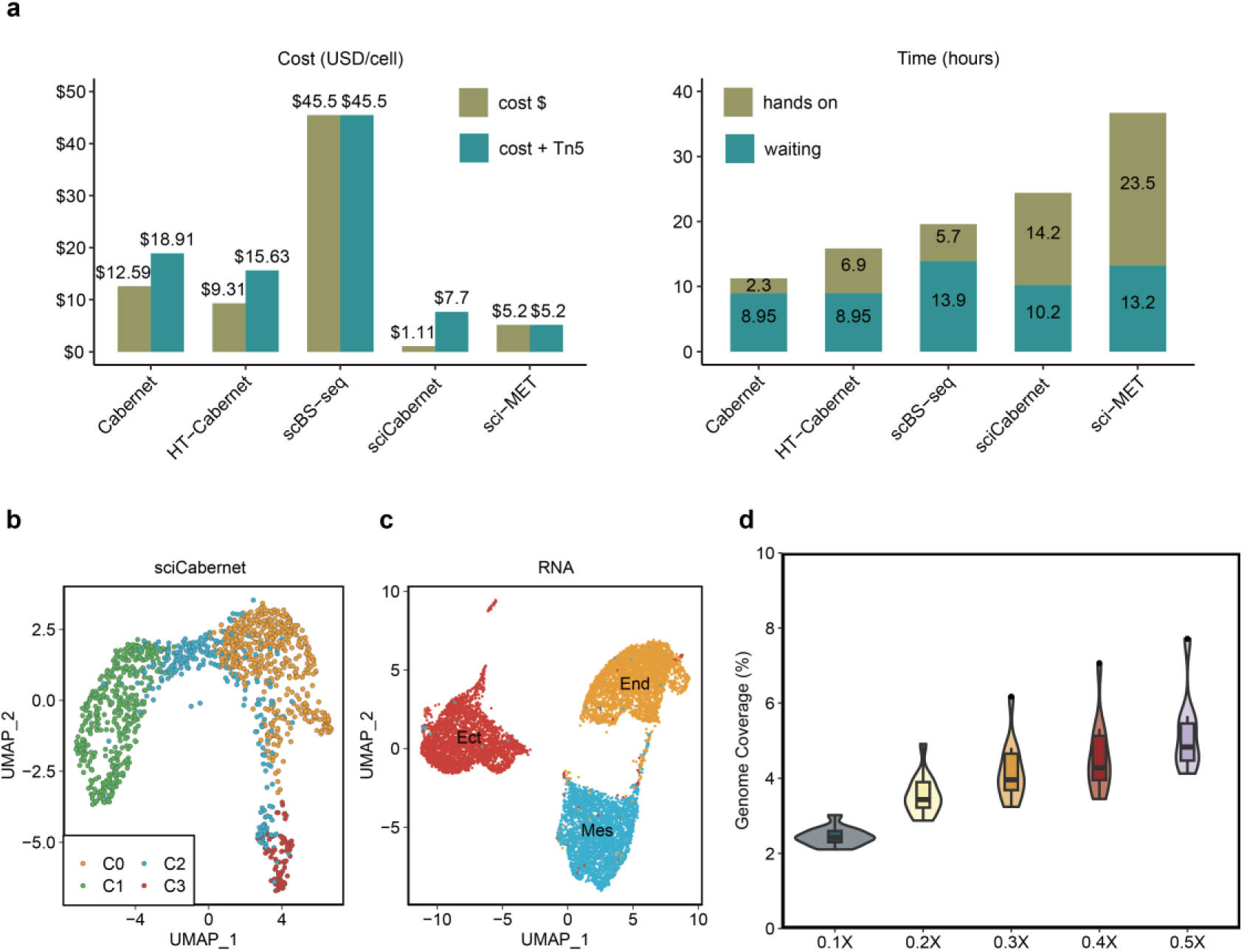
Cost-effectiveness and performance of Cabernet technology. (**a**) Cost of money and time by different sequencing methods. The yellow bar refers to the cost of reagents; the green bar refers to the combined cost of reagents plus Tn5. Detailed calculations of cost are shown in Table S6. **(b)** UMAP showing the clustering of E7.5 mouse embryo cells sequenced by sci-Cabernet. **(c)** UMAP showing the clustering of E7.5 mouse embryo cells based on 10x scRNA-seq data. **(d)** Violin plot showing the genome coverage of sci-Cabernet at different sequencing depths in K562 cells.

